# Genomic Insights into Hybridization and Speciation of Mitten Crabs in the *Eriocheir* Genus

**DOI:** 10.1101/2024.12.10.627874

**Authors:** Jun Wang, Xin Hou, Xiaowen Chen, Roland Nathan Mandal, Nusrat Hasan Kanika, Chunhong Yuan, Yongju Luo, Chenghui Wang

**Author notes:** Equal contribution (Jun Wang, Xin Hou, Xiaowen Chen).

## Abstract

Hybridization is a prominent and influential phenomenon with significant implications for adaptive evolution, species distribution and biodiversity. However, the intricacies of how hybridization influences genomic structure and facilitates adaptive evolution still elude our full understanding. Analyzing whole genome data from seven populations within *Eriocheir* genus across diverse geographic regions, we have validated a complex hybridization history between Chinese and Japanese mitten crabs. This hybridization gave rise to two distinct ecological species: Hepu and Russian mitten crabs with unique genomic architectures and adaptations. Genes related to reproduction, development, and temperature adaptation exhibit divergent selection signals, potentially contributing to their phenotypic diversity and ecological niches. Meanwhile, genes associated with reproduction, namely *Birc6*, *Bap31*, and *Poxn*, displayed robust evidence of selective sweeps in Hepu mitten crab. Notably, the favored alleles for these genes originated from the parental lineages during the hybridization process. Furthermore, Hepu mitten crab is a homoploid hybrid species that originated from an ancient hybridization event, resolving its longstanding taxonomic controversy. Our study sheds light on the evolutionary history of mitten crabs and highlights the crucial role of hybridization in driving adaptation, range expansion, and diversification within the *Eriochier* genus.

## Introduction

Environmental factors such as climate, water chemistry, and food resources often exert strong selection pressure on environmental adaptation and constrain the geographic distributions of species [1–5]. Hybridization between species, subspecies, or diverged populations serves as a potent driver of increased genetic diversity, reduced genomic vulnerability, and accelerated adaptive evolution, promoting both speciation and radiation [6–12]. Increased genetic diversity through hybridization can promote the natural geographic range expansion and novel genomic architectures in hybrids [13]. However, the full extent of hybridization’s role in global species diversity and the specific contribution of admixed genomes to genomic architecture for adaptive evolution remain subjects of debate and ongoing research [14–16].

It is evident that hybridization and introgression in hybrid zones promote phenotypic and genotypic adaptations [14,16]. This process is considered to be a potent force for speciation [17–19]. After hybridization, certain genes may be selected, introducing advantageous alien alleles merging into the genetic backgrounds of the parental populations. This process can consequently facilitate geographical expansion and promote adaptive radiation [20–22]. Numerous empirical studies across various animal groups, including butterflies, fishes, birds, and mammals, have substantiated the prevalence of homoploid hybrid speciation [18,19,23–26], suggesting that the increase of species diversity through hybrid speciation may be widespread and hence more imperative for the global species radiation [19,27].

Taxonomy of *Eriocheir* species, comprising *Eriocheir sinensis* (H. Milne Edwards, 1854) (Chinese mitten crab), *Eriocheir japonica* (De Haan, 1835) (Japanese mitten crab), and *Eriocheir hepuensis* (Dai, 1991) (Hepu mitten crab) has long been controversial due to their morphological similarities and limited molecular evidence [28–30]. Especially in the case of *E. hepuensis*, there is ongoing debate regarding whether it should be classified as a subspecies of Japanese mitten crab, an independent species, or a hybrid [30–36]. While, *E. sinensis* and *E.japonica* have long been considered as two independent species [31,36]. Mitten crabs have native ranges in East Asia: *E. sinensis* is mainly distributed in the Liao River, Yellow River, and Yangtze River from the northern to the middle part of China [35]. *E. japonica* inhabits the northern East Coast of Korea and Japan [33,37], and *E. hepuensis* is limited to the Nanliu River in southern China [28]. Additionally, invasive mitten crabs species, *E. sinensis* and *E. japonica* have been reported in Europe and North America [30,38–41], while *E. hepuensis* has been established in Iraq and Iran [42,43]. However, due to the difficulty in classifying the *Eriocheir* species only by subjective morphology characters, the exact invasive species of *Eriocheir* in non-native regions remains obscure [30,42]. Recent studies suggest that invasive *Eriocheir* species in Europe may be hybrids [41,44] and previously identified *E. sinensis* in European waters were molecularly determined to be *E. japonica* [41], underscoring the need to establish the taxonomic status of the *Eriocheir* genus and clarify the origin of *E. hepuensis* through robust genome data.

*E. sinensis* is an indigenous species in China and one of the most economically significant freshwater species, supporting a vast aquaculture industry in the country. It is widely cultured in the Yangtze river region, whereas *E.japonica* and *E.hepuensis* have not been cultivated as extensively [28]. Previous studies reported that the natural hybridization events occurred between *E. sinensis* and *E. japonica*. Two hybrid zones exist between *E. sinensis* and *E. japonica* in Vladivostok, Russia, and the Min River, China [35,45]. Meanwhile, research suggests that *E. hepuensis* in Nanliu River (Hepu, China), may also be a hybrid [30]. Hybridization events across varying latitudes result in distinct ecological hybrids that inhabit different ecological niches (southern versus northern), although they have the same parental species (*E. sinensis* and *E. japonica*). This fascinating mitten crab system gives us a unique opportunity to investigate the influence of interspecies hybridization on local environmental adaptation and ecological distribution.

In this study, we employ our previously published chromosome-level reference genome of *E. sinensis* [46] and combine the two *de novo* assembled draft genomes of *E. japonica* and *E. hepuensis* to delve into the intricate world of mitten crab hybridization. Our research aims to investigate the genome characteristics, population structure, genetic variations, and hybridization history of seven mitten crab (*Eriocheir* genus) populations across East Asia. Specifically, we seek to explore how gene flow between representative Chinese, Japanese, Russian, and Hepu mitten crabs has led to new speciation events, driven by genetic mixing. By focusing on physiological and phenotypic differences among these populations, we aim to assess how hybridization contributes to environmental adaptation and speciation. Furthermore, we intend to elucidate the genetic mechanisms behind hybrid genome evolution and how they influence adaptation to distinct ecological environments and distribution patterns.

## Results

### Genome assembly and annotation comparison of *Eriocheir* species

The high-quality chromosome-level genome of *E. sinensis* was assembled in our previous study [46]. The genomes of *E. japonica* and *E. hepuensis* were sequenced and *de novo* assembled in this study (**Table S1, Table S2**). The assembled genome sizes were 1.24 Gb for *E. japonica* and 1.18 Gb for *E. hepuensis*, with N50 lengths of 442,749 bp and 122,458 bp, respectively (**Table S3**). Using prediction evidence and RNA sequencing data, 18,418 protein-coding genes were identified in *E. japonica* and 19,253 in *E. hepuensis* which is comparable with *E. sinensis* (20,286 protein-coding genes) (**Table S3, Table S4**).

### Geographical and morphological diversity in mitten crab populations

Mitten crab samples were collected from seven geographic locations across different latitude, which is Chinese mitten crab collected from Liao River (LR, China), Yellow River (YeR, China), Yangtze River (YaR, China), and Min River (MR, China); Russian mitten crab collected from Vladivostok (VL, Russia); Japanese mitten crab collected from Hokkaido (HO, Japan), and Hepu mitten crab collected from Hepu (HP, China) (**Figure 1A, Table S5**). Among these populations, four exhibited distinct morphological differences in body weight and shape (**Figure 1B, 1C, Figure S1**). As shown in **Figure 1B**, the Hepu-HP mitten crab population has a notably smaller carapace and body size than the others. Chinese-LR, YeR, YaR mitten crab have nearly circular carapaces with deeper frontal teeth (A1), conspicuous lateral teeth, greater carapace width (A6), and longer limbs (F1, F2) (**Figure 1C**, **Figure S1**). In contrast, Japanese-HO mitten crab has nearly square carapaces with blunt frontal teeth (A1), less prominent lateral teeth, narrower carapace width (A6), and shorter limbs (F1, F2) (**Figure 1B and Figure S1**).

**Figure 1.**
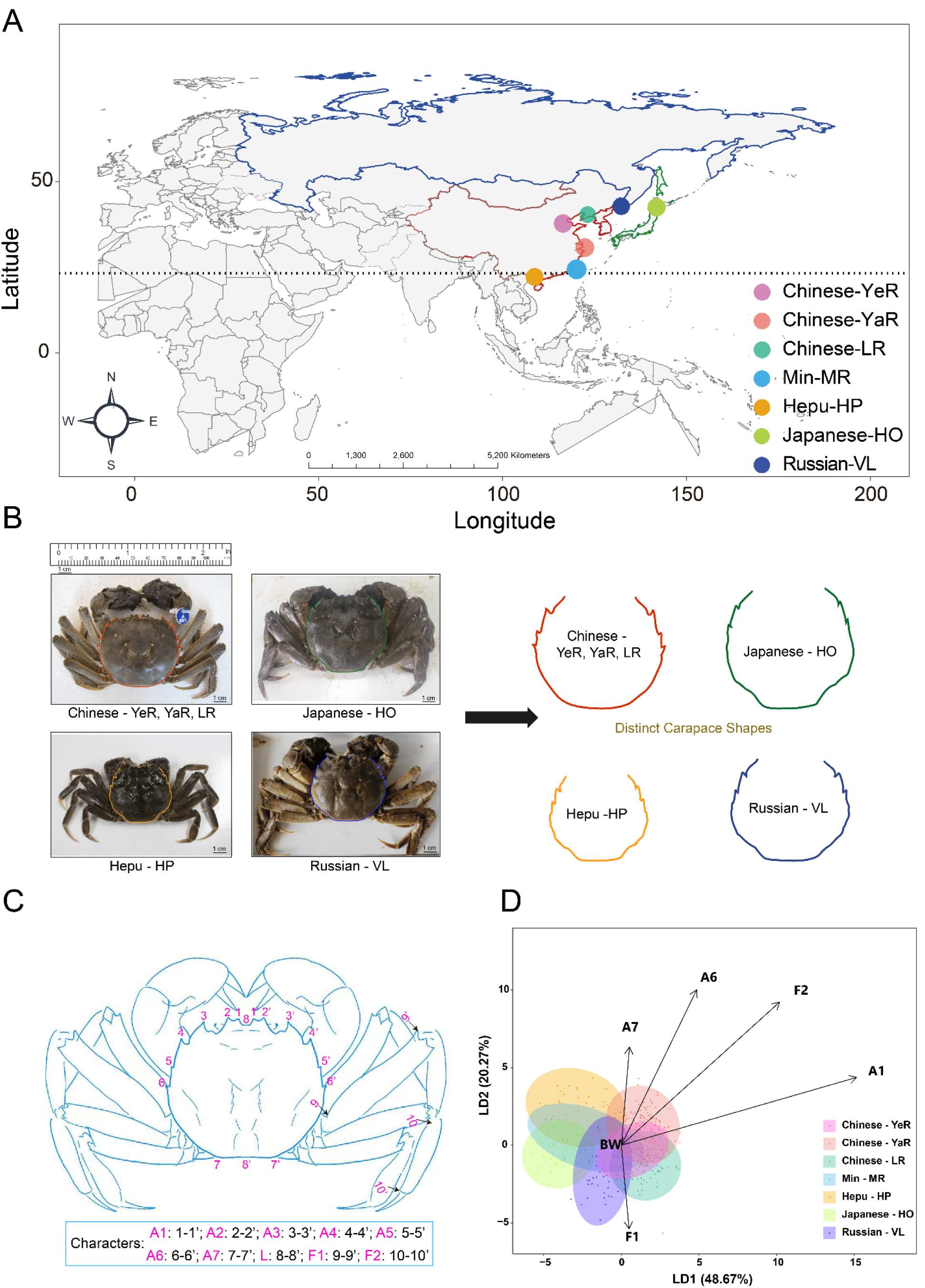
The Distribution and Phenotypic Characteristics of Mitten Crabs. (**A**) Depicts the geographical distribution of seven collected mitten crab populations, with their respective locations marked by circles. (**B)** Features images of mitten crabs. (**C**) Illustrates various measurement points used to calculate the external characteristics of mitten crab population. (**D**) Linear Discriminating Analysis (LDA) based on phenotypic measurements of mitten crabs collected from seven populations.

The ANOVA results further showed that southern populations, Hepu-HP and Min-MR mitten crabs have the smallest adult body weight **(Figure S1)**. In the Japanese-HO mitten crab, width of first pair of frontal teeth (A1), carapace width (A6), femur length (F1), and tibia length (F2) were the smallest, while these traits were the largest in the Chinese mitten crab (**Figure S1**). As for Hepu-HP and Russian-VL mitten crabs, we identified A1, A6, F1, and F2 to show intermediate levels compared with Chinese-LR, YeR, YaR and Japanese-HO mitten crabs, which were consistent with previous research (**Figure S1**) [45]. However, Russian-VL mitten crab presents a larger body weight, and smaller A1 and A6 compared with Hepu-HP mitten crab (**Figure S1**).

Additionally, Linear Discriminant Analysis (LDA) revealed distinct phenotypic clusters among the seven populations, with the first linear discriminant component (LD1) separating Chinese-LR, YeR, YaR from Japanese-HO mitten crabs, indicating distinct phenotype variation (**Figure 1D**).

Moreover, the time required for sexual maturation varies among the mitten crab populations from different regions **(Figure S2A)**. Hepu mitten crab, located in a high-temperature zone, reach sexual maturity within one year. In contrast, Japanese and Chinese mitten crab require two years, while Russian mitten crab, found in a lower-temperature zone, take three years to mature, reflecting the influence of regional temperature on their maturation periods **(Figure S2)**.

### Ancestral history of populations within the *Eriocheir* genus

We conducted genome resequencing and variant discovery based on 139 samples of mitten crabs at the whole genome level using *E. sinensis* chromosomal-level genome as reference [46]. We obtained a total of 5,547.87 Gb of raw data, with 39 Gb (around 23-fold depth) on average for each individual (**Table S6**). After quality filtering, all the sequenced reads were mapped to the *E. sinensis* reference genome, the average mapping rate was above 97% (**Table S6**). We identified a total of 10.27 million high-quality SNPs after filtration (**Table S7**).

Principal Component Analysis (PCA) analysis of 912,659 SNPs (Minor Allele Frequency > 0.05 and linkage disequilibrium pruned) showed clear genetic clustering among mitten crab populations. Four major clusters were observed, corresponding to the Chinese-YeR, YaR, and LR, Japanese-HO, Hepu-HP, and Russian-VL populations. Additionally, the Min-MR population appeared in an intermediate position between the northern and southern clusters **(Figure 2A)**. Meanwhile, a maximum-likelihood phylogenetic tree constructed from the whole-genome sequence data also confirmed the four clear clusters, with YaR, YeR, and LR clustering together representing the Chinese mitten crab lineage. Hepu-HP, Russian-VL, and Japanese-HO mitten crabs cluster together to represent the Japanese mitten crab lineage. Min-MR population shows genetic similarities to the Chinese mitten crab, with several individuals clustering closely, suggesting potential admixture (**Figure 2B**).

**Figure 2.**
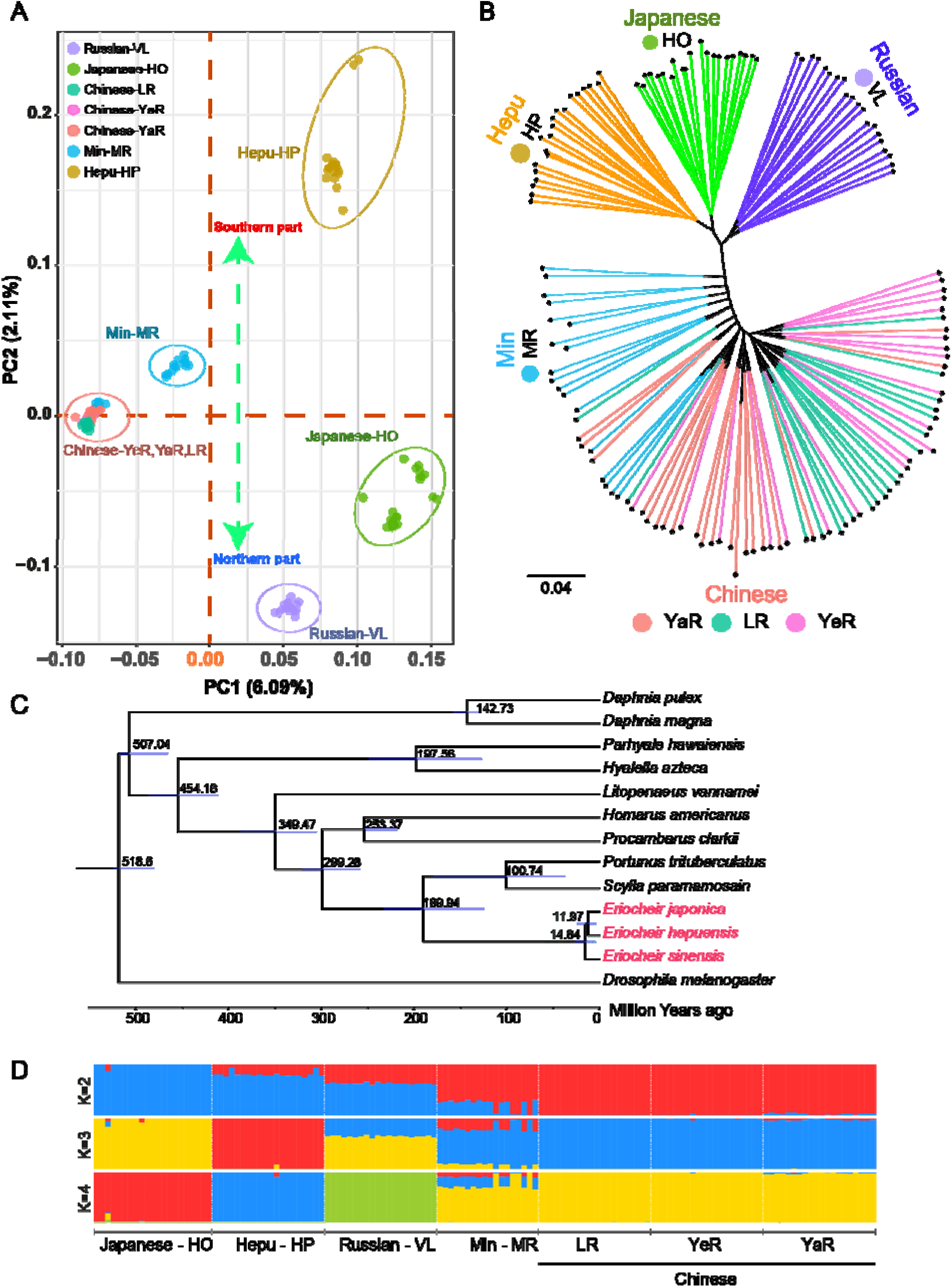
Genomic Insights into Mitten Crab Population Structure and Evolutionary History on 7 Mitten Crab Populations. (**A**) Genome-wide SNP-based Principal Component Analysis (PCA). (**B)** Phylogenetic tree constructed from high-quality SNPs, reveal the evolutionary relationships among those populations. (**C**) A phylogenetic tree was generated from one-to-one orthologs of 12 crustacean species with *D. melanogaster* as the root; estimated divergence times are provided beneath the tree. (**D**) Population structure analysis of the different mitten crab populations, considering K values ranging from 2 to 4.

In addition, comparative genomic analysis of 12 crustacean species and *Drosophila melanogaster* as an outgroup identified 684 one-to-one single-copy orthologous genes for phylogenetic assessment. The maximum-likelihood phylogenetic tree and molecular dating analysis revealed that *E. sinensis* (Chinese-YaR, YeR, and LR) and *E. japonica* (Japanese-HO) represent distinct evolutionary lineages, diverging approximately 14.84 million years ago **(Figure 2C)**. This divergence supports their classification as separate species within the *Eriocheir* genus. Moreover, phylogeny analysis revealed a close clustering of *E. japonica* and *E. hepuensis*, indicating a strong phylogenetic relationship between these species **(Figure 2C).** Additionally, a phylogenetic tree based on mitochondrial genomes indicates a close relationship between *E. sinensis* and *E. hepuensis* (Hepu-HP) (**Figure S3A**). The contrasting results from nuclear and mitochondrial genome analyses suggest historical hybridization events may have influenced the genetic makeup of the Hepu-HP mitten crab (**Figure S3A**). Together, these findings confirm that the Chinese-YaR, YeR, LR and Japanese-HO mitten crabs are genetically distinct species, despite evidence of occasional genetic exchange in related populations.

Further, admixture analysis revealed distinct yet interconnected populations within the *Eriocheir* genus. At *K* =2, there is a clear separation between the Chinese-YaR, YeR, LR and the Japanese-HO mitten crab populations, with admixture signals detected in Hepu-HP, Russian-VL, and Min-MR populations. At *K*=3, the Hepu-HP mitten crab population aligns separately with Chinese -YaR, YeR, LR and Japanese-HO mitten crab, with mixed ancestry is evident in the Russian-VL and Min-MR populations. At *K*=4, further division shows that Min-MR is a hybrid with contributions from Chinese-YaR, YeR, LR, Japanese-HO, and Hepu-HP lineages (**Figure 2D**). Results from Treemix analysis support these findings, indicating significant gene flow from the Chinese -YaR, YeR, LR mitten crab populations to the Hepu-HP, Russian-VL, and Min-MR populations, signifying extensive hybridization and introgression within the *Eriocheir* genus (**Figure S3B**). Pairwise sequentially Markovian coalescent (PSMC) analysis indicated small effective population size in Japanese-HO mitten crab (**Figure S3C**).

### Genomic differentiation and selection in *Eriocheir* genus

Analysis of nucleotide diversity (Pi) among biallelic SNPs in mitten crab populations revealed substantial differences between the Chinese-YaR, YeR, and LR and Japanese-HO lineages. The Chinese-YaR, YeR, and LR populations exhibited the highest nucleotide diversity (*Pi* = 3.59×10 ³), in contrast to the Japanese-HO population, which had the lowest *Pi* value at 3.05×10 ³ **(Figure 3A).** This reduction in genetic diversity within the Japanese-HO population suggests a potentially restricted gene flow over time.

**Figure 3.**
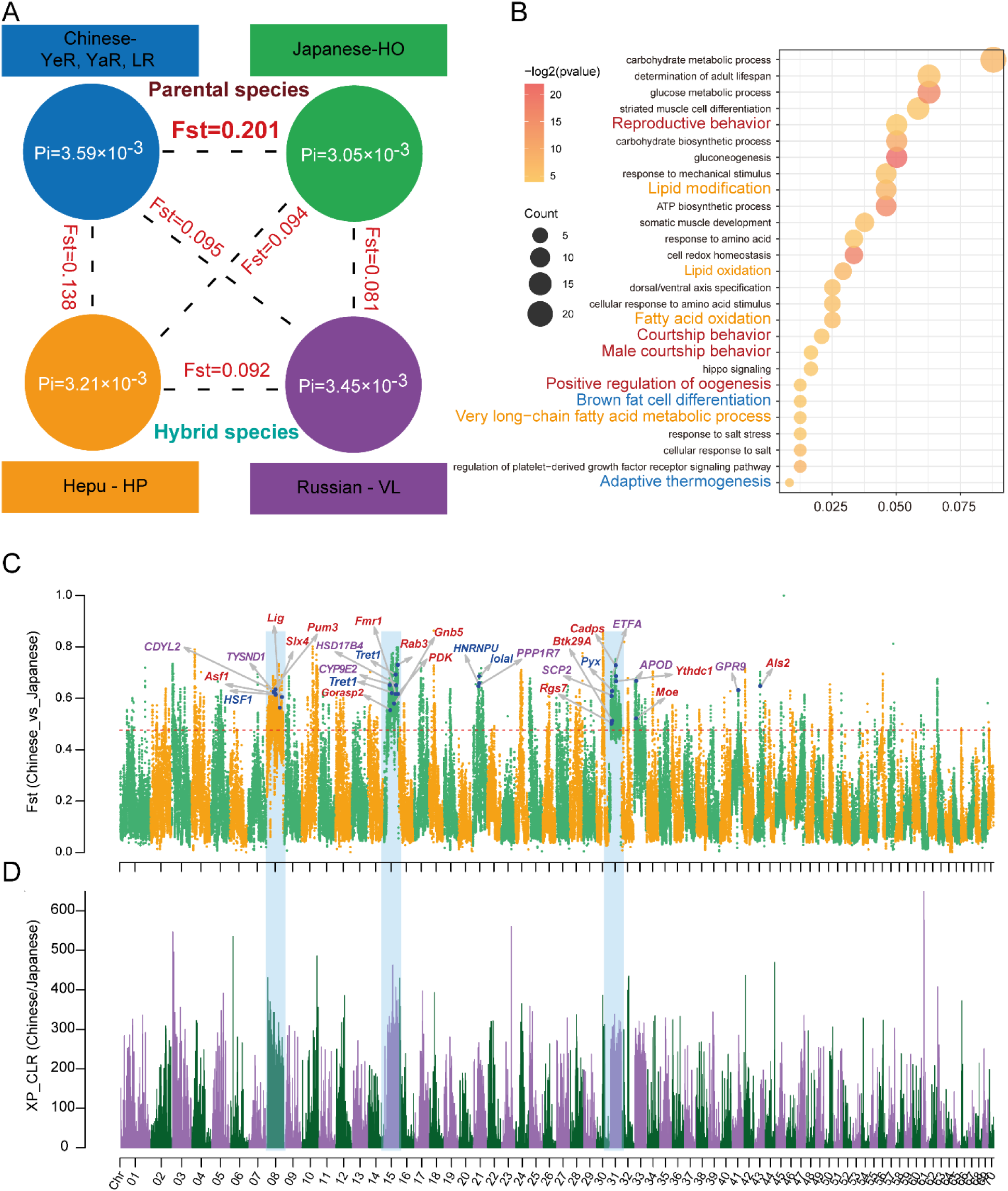
Genetic Differentiation Between Chinese and Japanese Mitten Crab in *Eriocheir* Genus. (**A**)Nucleotide diversity (Pi) within population and genetic differentiation index (*Fst*) between two populations. (**B)** GO enrichment analysis of genes that showed divergent selection between Chinese and Japanese mitten crab. (**C)** Genomic-wide depiction of genetic differentiation (*Fst* values) between Chinese and Japanese mitten crabs. Genes linked to the reproductive process, lipid oxidation, and temperature stimulus are marked with red, purple, and blue colors. (**D)** Genome-wide visualization of selection signals (XP-CLR scores) between Chinese and Japanese mitten crabs.

Whole-genome analyses of genetic differentiation further underscored the substantial evolutionary distance between the Chinese and Japanese mitten crab populations. The *Fst* value, which measures population differentiation, revealed the largest genetic differentiation between the Chinese-YaR, YeR, LR, and Japanese-HO populations, marked by a mean weighted *Fst* value of 0.201 (**Figure 3A**). In contrast, the smallest genetic differentiation was observed among the three populations (LR, YeR, and YaR) of the Chinese mitten crab (**Figure S4**). Moreover, largest average *Dxy* values were identified between Chinese-LR, YeR, YaR and Japanese-HO mitten crab populations (**Table S8**).

We identified notably differentiated genomic regions when comparing Chinese-YaR, YeR, LR and Japanese-HO mitten crabs, comprising the top 1% based on *F*st values. These regions encompassed a total of 401 genes (**Table S9**). GO enrichment analysis indicated that these genes were enriched in lipid oxidation, muscle cell differentiation, determination of adult lifespan, adaptive thermogenesis, courtship behavior, and so on (**Figure 3B**). Highly genomic differentiation regions were identified in almost the entire chromosomes, such as Chr 08, Chr15, and Chr 31 (mean *Fst* value above 0.6), these findings suggest that the Japanese-HO and Chinese-YaR, YeR, LR mitten crabs may have undergone divergent selection in these specific genomic regions after their speciation (**Figure 3, C and D**). Functionally involved genes such as *Asf1*, *Lig*, *Btk29A*, *Slx4*, *Als2* associated with reproduction process, *Tysnd1*, *Cyp9e2*, *Hsd17b4*, *Apod* associated with lipid oxidation, and *Hsf1, Tret1, Pyx* associated with temperature stimulus showed strong genetic differentiation and selection signals between Chinese and Japanese mitten crab (**Figure 3, C and D**).

Previous studies have identified the *Btk29A* gene, known for its role in courtship behavior and male genitalia development, as showing selection signals within the Japanese-HO mitten crab population [47,48]. Two distinct haplotypes of the *Btk29A* gene were identified in the Japanese-HO and Chinese-YaR, YeR, LR mitten crab populations. A single nucleotide mutation on Chromosome 31 (position 3522309, G to A), led to alternative splicing in the *Btk29A* gene, causing the Japanese-HO mitten crab to lose exon 2 (**Figure S5, A and B**). Besides, the expression of the *Btk29A* gene exhibits significant difference between Chinese-YaR and Japanese-HO mitten crabs in the ovaries **(Figure S5 C)**. Additionally, the *Lingerer* (*Lig*) gene known for its role in copulation initiation and termination [49] exhibited signs of divergent selection signals between the Japanese -HO and Chinese -YaR, YeR, LR mitten crab populations.

A nucleotide change from A to C at position 1891 caused an amino acid substitution from methionine to leucine (Met631Leu), resulting in structural alternations in the lingerer protein (**Figure S5, C and D**). Also, the expression of the Lingerer (*Lig*) gene differs between Chinese-YaR and Japanese-HO mitten crabs (**Figure S5 F**).

Simultaneously, the *Pyx* gene, responsible for cation channel activity linked to protection and tolerance against high-temperature stress [50], and the *Hsf1* gene, which plays a role in the heat shock response, displayed significant divergence with two distinct haplotypes identified between the Chinese-LR, YeR, YaR and Japanese-HO mitten crabs (**Figure S6**). Furthermore, the expression of both *Hsf1* and *Pyx* genes showed notable differences between Chinese-YaR and Japanese-HO mitten crabs (**Figure S6**). Notably, missense mutations, specifically (c.1924 G>C) in the *Pyx* gene and (c.732A>G) in the *Hsf1* gene, resulted in amino acid changes from Asp to His and Ile to Met in the Japanese-HO mitten crab (**Figure S6**). Additionally, Hepu-HP and Russian-VL mitten crabs, both of which are hybrids, exhibited a substantial proportion of heterozygous genotypes within their populations (**Figure S6**).

### Different genome architecture of the two hybrid mitten crabs

Our results indicated that the Hepu-HP and Russian-VL mitten crabs have admixture ancestry (**Figure 2**). In both hybrid populations, there is a predominant genetic inheritance from the Japanese mitten crab, along with a lesser contribution from the Chinese mitten crab. In terms of their genome structure, Chinese and Japanese mitten crabs contribute about 19.59% and 80.41% to the genomic composition of Hepu mitten crabs, and 37.87% and 62.13% to that of Russian mitten crabs, respectively (**Figure 4A**). Interestingly, these two hybrid populations occupy different ecological environments at varying latitudes and display significant genomic differentiation across their entire genomes (**Figure 4C**). Identified, 1734 highly differentiated genomic regions between the two hybrid populations (Top 1% regions of *Fst* value), identified 343 regions in Chromosome 06 and 203 regions in Chromosome 70, and no significant differentiated regions were identified in Chromosome 08 and Chromosome 31 like their parental lineages, indicating Hepu-HP and Russian -VL mitten crab did not share the differentiated regions with their parental species (**Figure 4C**). Furthermore, among the 390 genes identified, there is a substantial genomic differentiation between the two hybrid populations. (**Figure 4, B and C, Table S10**). In summary, the results of the GO enrichment analysis reveal that these genes are enriched in functions related to reproductive behavior, autophagy, muscle cell differentiation, and the regulation of ovulation, among others (**Figure 4B**). These findings suggest hybrid populations may have experienced divergent selection following hybridization in distinct ecological environments.

**Figure 4.**
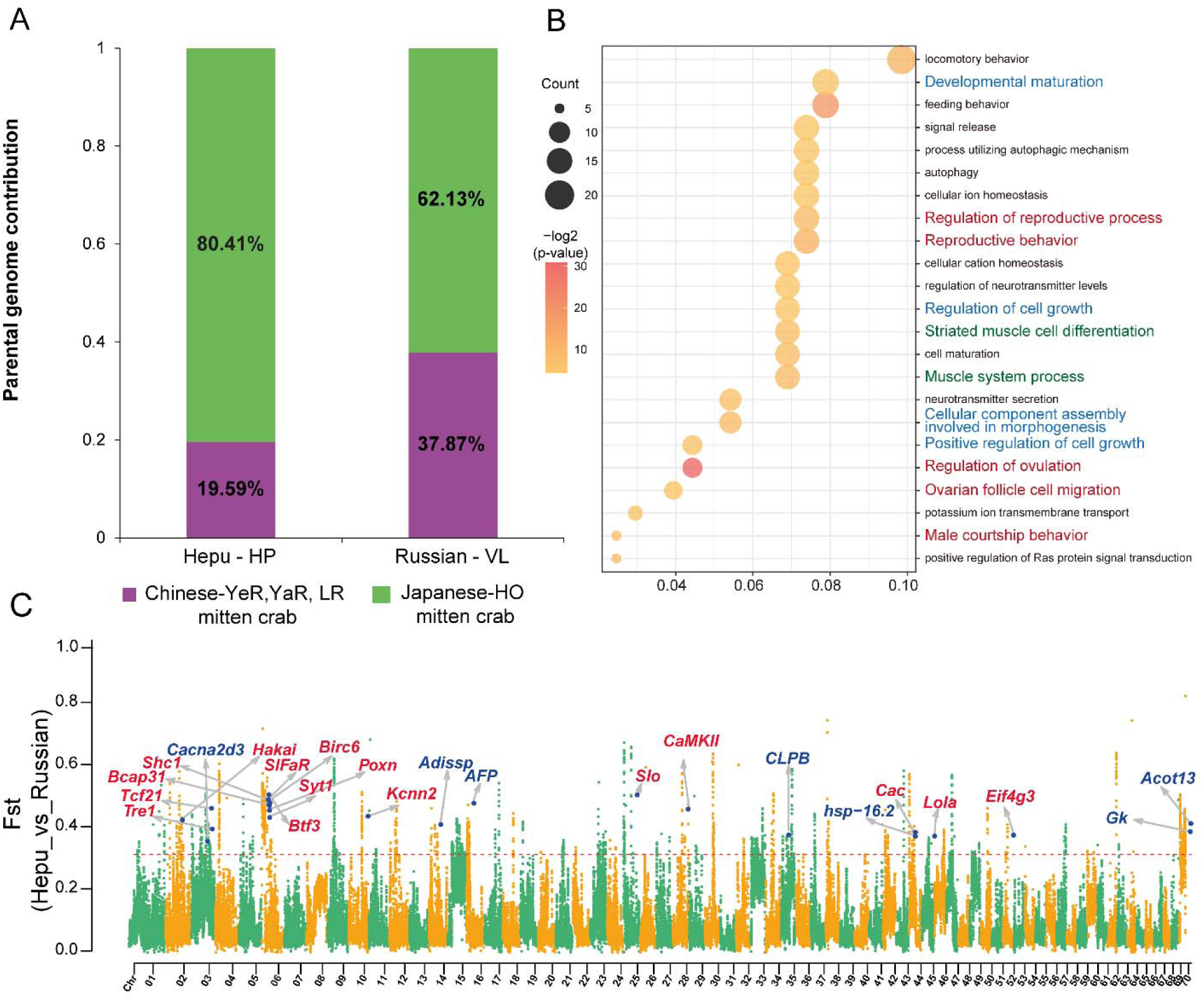
Different Genome Architecture Between Hepu-HP and Russian-VL Mitten Crab. (**A**)Illustration of parental genome contributions. (**B)** GO enrichment analysis of genes exhibiting genetic differentiation. (**C)** A genome-wide representation of genetic differentiation (*Fst* values) with genes associated with the reproductive process and temperature stimulus marked in red and blue

The mean *F*st values were 0.138 and 0.094 between Hepu mitten crab and their parents, and 0.095 and 0.081 between Russian mitten crab and their parents (**Figure 3A, Figure S7**). Moreover, the *F*st value between the two hybrid mitten crabs was 0.092 (**Figure 3A**). These findings suggest that both Hepu and Russian mitten crabs possess distinct genome architectures and are undergoing an intermediate level of genetic differentiation (**Figure 4C**). Of the divergent SNPs (fixed) identified between the Chinese and Japanese mitten crabs, a notable majority, comprising 87.65% and 98.84% of the SNPs, showed heterozygous genotypes in Hepu and Russian mitten crabs (**Figure S8**). Additionally, 1.17% and 0.46% of the SNPs represented the Chinese mitten crab genotype, while 11.18% and 0.70% displayed the Japanese mitten crab genotype in Hepu and Russian mitten crabs, respectively (**Figure S8**).

### Natural Selection in Hepu mitten crab

Remarkably, we detected robust signals of selective sweeps in Chromosome 06 specific to the Hepu-HP mitten crab (**Figure 5A**). Within this region, numerous genes, including *Btf3, Birc6, Poxn, Syt1,* and *Shc1,* displayed clear indications of natural selection acting on the Hepu mitten crab genome. These selected genes are associated with the reproductive process and present nonsynonymous mutations in the Hepu mitten crab indicating that these reproductive-related genes in Chromosome 06 were selected by the natural environment and may affect the reproductive isolation (RI) (**Figure A, B, Figure S9**). Notably, the homozygous mutations were predominantly exclusive to the Hepu-HP mitten crab, while only reference allele genotypes were present in the Chinese mitten crab populations (LR, YeR, and YaR). Considering the Chinese mitten crab genotype as reference, the Japanese -HO mitten crab population exhibited only a small proportion of homozygous mutations (**Figure 5B**). The homozygous mutations observed in the Hepu mitten crab, likely influenced by strong selection, may have originated from the Japanese mitten crab through a hybridization event.

**Figure 5.**
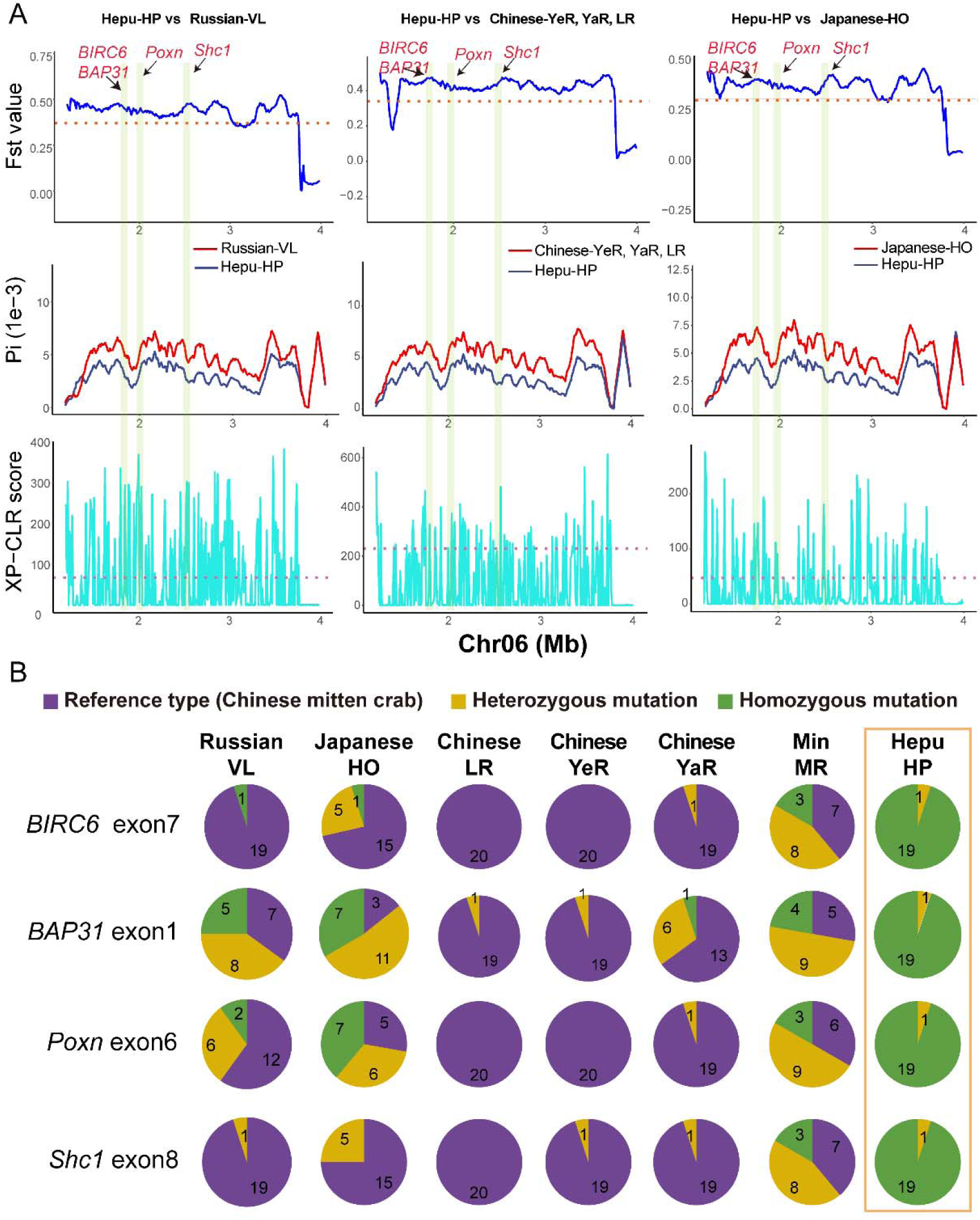
Detection of Selection Signals in Hepu Mitten Crab. (A) Genomic comparisons of population differentiation (*Fst*), nucleotide diversity (*Pi*), and XP-CLR signals across mitten crab populations in specific regions of Chromosome 6. Highlighted genomic regions include key loci such as *BIRC6, BAP31, Pxn,* and *Shc1*, which show signals of selection in Hepu mitten crab. (B) Pie charts summarizing mutation profiles across populations, highlighting proportions of reference types (purple), heterozygous mutations (yellow), and homozygous mutations (green), alongside the frequency distribution of distinct gene haplotypes. Number in the circle indicates the number of individuals in this population with the specific genotype.

Moreover, we also identified several genes, such as *AFP* and *Hsp-16.2*, associated with temperature stimulus, displaying selection signals in the Hepu mitten crab. Notably, nonsynonymous mutations (c.338 C>T) occurred in the *AFP* gene, while (c.269 C>G) occurred in the *Hsp-16.2* gene, resulting in amino acid changes from Pro to Leu and Thr to Ser in the Hepu mitten crab (**Figure S10, A and B**). These genetic variations may have implications for temperature adaptation differences between the Hepu-HP mitten crab and Russian-VL mitten crab.

## Discussion

The unique central distribution of Chinese-LR, YeR, YaR and Japanese-HO mitten crabs have driven the formation of two ecologically significant hybrid populations, Russian-VL mitten crab to the north and Hepu-HP mitten crab to the south, demonstrating how geographical proximity and genetic intermixing foster adaptive diversity within the *Eriocheir* genus (**Figure 6**) [45]. Genetic and phenotypic analyses suggest that the Hepu-HP mitten crab has diverged significantly, potentially representing a distinct species within the *Eriocheir* genus. Through our research, we have illuminated the intricate history of hybridization within the *Eriocheir* genus. This discovery resolves long-standing taxonomic debates that have spanned decades [30,31,33].

**Figure 6.**
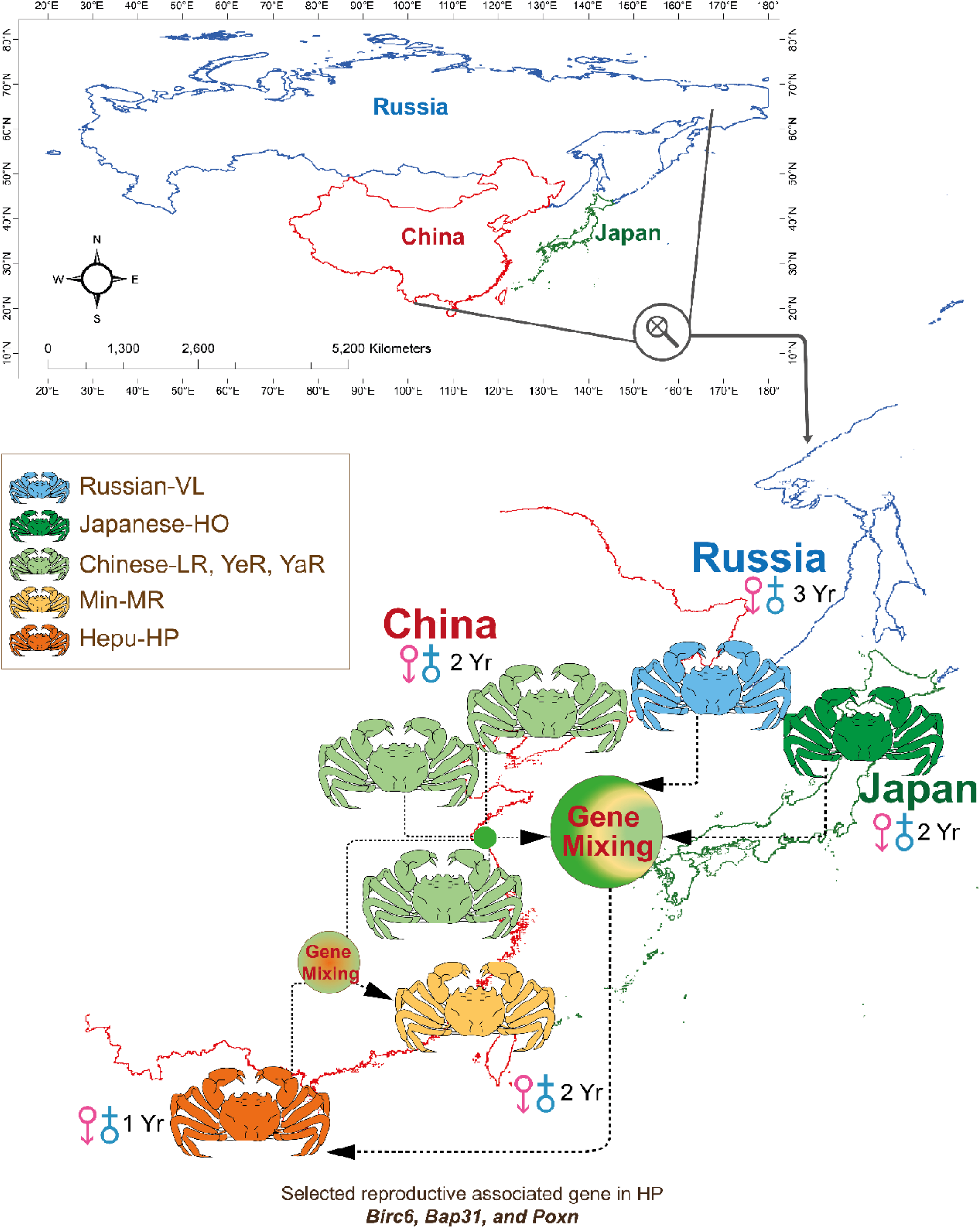
Depicts of Hybridization and Ecological Distribution of *Eriocheir* genus in East Asian region. Figure illustrates the genetic hybridization patterns and distribution of mitten crabs across East Asia. The Hepu mitten crab exhibit genetic mixing with Chinese and Japanese mitten crabs, acting as a hub of gene exchange. Additionally, the Min (MR) crab population is shown to mix genetically with both Hepu and Chinese mitten crab, further contributing to regional genetic diversity. Arrows indicate the direction of gene flow, while colored circles represent areas of genetic mixing.

### Multilayered hybridization history of the *Eriocheir* genus

The Chinese-YeR, YaR, LR, Japanese-HO, Russian-VL, and Hepu-HP mitten crab populations show distinct geographic, morphological, evolutionary, and genetic differences. Hepu-HP and Russian-VL mitten crabs originated from hybridization between Chinese and Japanese mitten crab, followed by divergent selection that led to distinctive phenotypic traits (**Figure 4, 6**), illustrating how hybridization and selection drive diversity within the *Eriocheir* genus. What’s even more intriguing is that mitten crabs collected from the Min River region exhibit genetic admixture signals, signifying their hybrid nature. These crabs display a complex admixture ancestry with genetic contributions from the Chinese, Japanese, and Hepu mitten crabs, reflecting their intermediate geographic position (**Figure 2D**, **Figure 6, Figure S3B**). This pattern suggests that hybridization in the Min-MR mitten crab likely occurred after the formation of the Hepu mitten crab, pointing to a more recent hybridization event involving Chinese and Hepu mitten crab [34,35]. In light of the intricate and recurring hybridization within the *Eriocheir* genus, it is reasonable to suspect that the invasive mitten crabs in Europe and North America may likewise be of hybrid origin, as earlier studies have proposed [44]. Confirming this hypothesis will necessitate additional research in the future.

Prior research has suggested that the Hepu mitten crab should be classified as an independent clade rather than a hybrid. This distinction can be attributed to two key factors: 1) The limited discriminatory power of utilizing only AFLP loci and COII markers, 2) the Hepu mitten crab is an ancient hybrid that has undergone adaptive evolution which will form unique genome architecture [51]. In this study, Hepu-HP mitten crab shows a high level of heterozygosity, likely due to its hybrid origin involving Chinese and Japanese mitten crab (**Figure S8**).

Reports suggest that Japan island separation from the southeastern margin of the Chinese continent 15-25 million years ago, driven by crustal movements [52,53], may contribute to the speciation event of the Chinese and the Japanese mitten crab. Mitten crabs are catadromous species that require seawater for reproduction. The isolation of the Japan sea from China’s marginal seas such as the Yellow sea and East sea, likely contributed to reproductive isolation and differentiation. This is because mitten crab, which are offshore spawners, cannot migrate to the deep sea to reproduce [35,51,54]. The divergence of the Chinese and Japanese mitten crab was estimated at around 14.84 million years ago, which aligns with the formation time of the Japan island (**Figure 2C**).

### Adaptive evolutionary path of Hepu-HP and Russian-VL mitten crabs through hybridization

The differences in morphological traits mirror the process of adaptive evolution, as seen in the unique ecotype phenotypes of the Russian-VL and Hepu-HP mitten crabs. These distinct characteristics provide potential evidence of their adaptations to diverse environmental conditions [45]. Diverse environments may impose strong selection pressure that has shaped their distinct phenotypes (**Figure 1, Figure S1**). Genomic analysis provides evidence that the Russian-VL and Hepu-HP mitten crabs have undergone divergent selection with specific genes related to their adaptation being selected (**Figure 4**). The observed differences in reproductive maturity and morphology between mitten crabs from northern and southern regions highlight how environmental factors shape species adaptations. Specifically, Hepu-HP mitten crabs, located in the warmer, southern region of China, mature within a year. In contrast, Russian-VL mitten crabs, residing in colder northern regions, take 3 years to reach maturity [55]. Analysis of Hepu mitten crab identified strong selected signals of genes related to their reproductive and temperature adaptation (**Figure 5**). The accelerated reproductive maturity in Hepu-HP mitten crabs may be an adaptive response to their warmer environment, which exceeds the annual mean temperature threshold (20) for optimal mitten crab growth and development [56]. The reported annual mean temperature in Hepu exceeding 20 may impose strong selection on the Hepu mitten crab. This adaptation may impose strong selective pressure on genes associated with reproductive timing and temperature resilience in the Hepu-HP population, contributing to their distinct reproductive strategies and morphology (**Figure S2**). We also observed that the body weight of the Russian-VL mitten crab is higher than that of the Hepu-HP mitten crab (**Figure S1**), which could suggest that environmental factors may play a role. A similar phenomenon was noted in mice, where northern population had significantly larger sizes compared to those closer to the equator [57]. It’s possible that Hepu (Guangxi, China) might not be the ideal habitat for mitten crabs. Nevertheless, hybridization could play a role in expanding the distribution of Hepu mitten crabs to southern areas, resulting in smaller individual sizes, improved heat tolerance, and varied sexual maturity times for better adaptation (**Figure 6**).

### Evidence for a hybrid speciation event

Despite naming his book “On the Origin of Species,” Darwin provides a sketchy blueprint for how species form in nature and therefore characterized this as the “mystery of mysteries.”[58]. A key part of hybrid speciation is how hybrids separate from their parent species by developing reproductive isolation (RI). This helps them keep their own genetic identity and evolve into a new species [22,59,60]. Research indicated that the reproduction season and sexual mature time (1 year) of Hepu-HP mitten crab is different from their parents [61]. In our research, we identified annotated reproductive-associated genes such as *Birc6, Bap31, Poxn, Syt1,* and *Shic1* which are involved in oogenesis, spermatogenesis, and courtship behavior showed strong selection signals in Hepu-HP mitten crab, suggesting that these genes may contribute to the development of reproductive isolation between the Hepu mitten crab and its parent species (**Figure 5**). However, the molecular function of these selected genes needs to be verified in future study. The Hepu-HP mitten crab is known for its abbreviated sexual maturity period, smaller individual size, and its unique habitat as the only mitten crab species living in the southernmost region with higher annual temperature (**Figure S2**). These factors account for the significant ecological differences observed between the Hepu-HP mitten crab and its parent species [28]. In the culture process of the Chinese mitten crab, a high temperature is not suitable for the survival of the Chinese mitten crab, and a high temperature will promote the precocity of the Chinese mitten crab, which is consistent with the characteristics of the Hepu mitten crab [61]. Together, our results pointed out that large genetic differentiation, significant phenotype variation, and reproductive isolation were identified between the Hepu mitten crab and its parental species, indicating the Hepu mitten crab is an independent ecological species formed through homoploid hybrid speciation event [22].

In conclusion, our study unravels a complex tapestry of hybridization within the *Eriocheir* genus. Notably, the Hepu-HP and Russian-VL mitten crabs, as key hybridized species, exhibit substantial genomic differentiation and have experienced adaptive evolution (**Figure 6**). To unlock the secrets of mitten crab adaptation, future research should investigate additional populations, including those in invasive regions, and explore the molecular function of the identified genes that have been selected by different environments. These intricate hybridization events have likely had a significant role in shaping species diversity within the *Eriocheir* genus and have had a significant impact on their global distribution patterns.

## Materials and Methods

### Sample collection and morphological characters measurement

A total of 139 individuals from 7 populations, Liao River (LR), Yellow River (YeR), Yangtze River (YaR), Vladivostok (VL), Hokkaido (HO), Min River (MR), and Hepu (HP), which include *E. sinensis*, *E. japonica*, *E. hepuensis*, and their hybrid populations were collected from 2015-2017 (**Figure 1A, Table S3**). The geographical location of all collected mitten crab individuals included in this study were recorded with a global positioning system on the WGS84 datum. All the collected adult mitten crab samples were measured for body weight (BW), shell length (L), shell width (A6), length of frontal teeth (A1-A5), and limb length (F1 and F2). All the morphological character data were normalized by dividing the shell length (L) except body weight. Linear Discriminate Analysis (LDA) analysis was conducted using the normalized morphological characters by machine learning methods and comparisons among the 7 populations for all the morphological characters were conducted by ANOVA analysis.

Temperature data from 1960 to 2020 was obtained from WorldClim version 2.1, a high-resolution climate database accessible at https://www.worldclim.org/data/worldclim21.html [62]. This dataset provides global, monthly temperature records at various spatial resolutions, supporting robust analysis of long-term climate trends. Our study, sampling procedures, and experimental protocols were approved by the Institutional Animal Care and Use Committee of Shanghai Ocean University (Shanghai, China) on the care and use of animals for scientific purposes.

### Genome assembly and Gene annotation of *E. japonica* and *E. hepuensis*

For *E. japonica*, an adult wild male individual from the Hokkaido was collected. Pair-end libraries of 180 bp, 500 bp, and 800 bp, as well as mate-pair-end libraries of 2 kb, 5 kb, and 10 kb, were constructed and sequenced on the Illumina HiSeq 2500 platform. Additionally, a 20 kb DNA library was created and sequenced on PacBio RS II platform. Furthermore, a 40 kb DNA library was constructed following the 10 X genomic sequencing library construction pipeline and sequenced on Illumina Hiseq 4000 platform. Initially, clean reads underwent assembly into contigs using Platanus 2.0.2 [63] and were subsequently linked via Redundans with default parameters [64]. Gaps were closed utilizing PBjelly2 with PacBio raw reads, followed by error correction [65]. Rascaf v1.0.2 was employed to further link the assemblies using transcriptome sequencing data [66]. Finally, scaffolds underwent further linked using 10X Genomics linking reads with ARCS v1.2.1 [67] and LINKS v2.0 [68].

For *E. hepuensis*, an adult wild male individual was collected. Pair-end libraries of 250 bp, 400 bp, and 800 bp, along with mate-pair-end libraries of 2 kb, 5 kb, and 10 kb, were constructed. These libraries underwent sequencing on an Illumina HiSeq 4000 platform. Subsequently, following the filtration of low-quality reads, all clean reads were assembled into contigs using SOAPdenovo2 with K=27, 31, and 99 [69]. The resulting contigs were further linked into scaffolds using Redundans with default parameters [64]. To refine the assemblies, Rascaf v1.0.2 was utilized for further linkage of the mitochondrial-genome-free assemblies, employing transcriptome sequencing data.

Five tissues (eyestalk, gill, hepatopancreas, muscle, and ovary) from *E. japonica* and *E. hepuensis* were collected and used for RNA-Seq sequencing. The RNA-Seq libraries were constructed with Truseq^TM^ RNA sample prep Kit for Illumina and sequenced on the Illumina HiSeq 4000 platform with pair-end mode (2X150bp). The RNA-seq data were used for genome assembly and annotation.

Transcriptome alignment, *de novo* gene prediction, and sequence homology-based predictions implemented in Evidence Modeler (EVM) were used for gene prediction [70]. Initially, RNA-Seq reads underwent transcriptome alignment and were assembled into transcripts using STAR-2.7.10a and Stringtie v2.2.0 [71,72]. These transcripts were then aligned to the genomes to acquire gene structure annotation information through PASA v2.5.1 [73]. For *de novo* gene prediction, GeneMark-ET and Augustus were used to predict genes on transposable-elements-hard-masked genome sequences [74,75]. A high-quality dataset was generated by PASA v2.5.1 to train these ab initio gene predictors. Additionally, sequence homology-based gene prediction incorporated protein sequences from the SwissProt vertebrates’ database and six organisms (*Eriocheir sinensis*, *Daphnia pulex*, *Lepeophtheirus salmonis*, *Caenorhabditis elegans*, *Drosophila melanogaster*, *Homo sapiens*) into GeMoMa v1.9 to generate homology gene structures. Subsequently, all predicted gene structures were integrated into consensus gene models using Evidence Modeler v2.0.0 [70]. To determine the functional annotation of the gene models, a BLASTP search was conducted against public protein databases, including NR (non-redundant protein sequences in NCBI), SwissProt, RefSeq, and KEGG with an E-value threshold of ≤ 1e-5. Additionally, the motifs and domains of each gene model were predicted using InterProScan 5 against public protein databases, including ProDom, PRINTS, Pfam, Gene3D, CCD, SMART, PANTHER, PROSITE and SUPERFAMILY [76].

### Whole genome resequencing libraries construction and resequencing

In each population, 18-21 individuals were selected for DNA extraction and sequencing librairy construction. DNA was extracted from muscle tissue by using the Qiagen Blood & Tissue DNEasy kit. The concentration and purity of extracted DNA were measured by NanoDrop 2000. After quality control, DNA sequencing libraries with 350-400 bp insert size were sequenced on the Illumina Novaseq 6000 platform with paired-end mode (150-bp read length).

### Genome read mapping and variant calling

Raw reads were filtered by Trimmomatic v0.39 using default parameters [77]. After filtering, all the clean reads from each individual were aligned to the *Eriocheir sinensis* reference genome using bwa 0.7.17 (parameter: -t 12 -M -R) [78]. The aligned bam files were then sorted using samtools 1.7 and PCR duplicates were marked by MarkDuplicates module by GATK 4.3.0.0 [79]. Then, the GATK HaplotypeCaller module was run on each bam file to generate gvcf (genomic variant call format) files. The GVCF files from the 139 individuals were combined into a single GVCF file by CombineGVCFs implemented in GATK 4.3.0.0, then the VCF files were obtained through GenotypeGVCFs command. The SNPs were further hard filtered using the following parameters: (1) QD < 2.0 || MQ < 40.0 || FS > 60.0 || SOR > 3.0 || MQRankSum < -12.5 || ReadPosRankSum < -8.0”, (2) read depths < 2, (3) only biallelic were kept. The SNPs were annotated by SnpEff v 5.0 software [80]. SNPs in coding sequences were classified as synonymous, nonsynonymous, intron, intergenic_region, and so on (**Table S7**).

### Population structure analysis

For the population structure and phylogenetic analysis, variants with a missing rate of >20% and a minor allele frequency (MAF) <0.05 were removed using Plink v1.9 [81]. Meanwhile, in order to avoid linkage-disequilibrium (LD) which will affect the population structure, all loci within 50 kb windows with an r2 exceeding 0.2 (--indep-pairwise 50 10 0.2) were filtered. The phylogenetic tree was constructed with IQtree v2.0.3 (-bb 1000 -mem 120G -nt 16 -m MFP+ASC-st DNA) with 1,000 bootstrap [82]. The population structure analysis was conducted using Admixture v2.34 with 30,000 burning and 20,000 interactions [83]. PCA analysis was conducted by smartpca version: 18140 using filtered biallelic SNPs. Gene flow were detected with Treemix v1.13 software, which infers a maximum likelihood tree based on genome-wide allele frequency data [84]. Demographic histories of the mitten crabs were reconstructed using the Pairwise Sequentially Markovian Coalescent (PSMC) model with the mutation rate (μ) was set to 7×10^-9^, and the generation time (g) was set to 2 years [85].

### Genome scans for natural selection

We used VCFtools 0.1.17 to calculate fixation index (*Fst*) for pairwise comparison of the 7 populations of mitten crabs with the biallelic SNPs [86]. The window size was set to 100 kb and the stepwise distance was 10 kb. The nucleotide diversity for each population was also calculated by VCFtools 0.1.17 with the window size set to 100kb and the stepwise distance was 10kb. To identify the regions under selection, we calculate the nucleotide diversity (Π), and genetic differentiation index (*Fst*) between populations with a sliding window of 100 kb and a 10 kb overlap between adjacent windows. Windows that were in the top 1% of *Fst* values and Π ratio were considered to be the putative selection target regions. In order to further verify the selection regions, the XP-CLR methods were also used to detect selective sweeps. Each chromosome was analyzed using the program xp-clr (--rrate 3.09e-8 --ld 0.95 --maxsnps 200 --size 100000 --step 10000)[87]. The windows in the top 1% of XP-CLR scores were considered as candidate candidate-selective regions.

### Mitochondrial genome assembly and Phylogenetic tree construction

The mitochondrial genome was *de novo* assembled by NOVOPlasty using the whole genome sequencing data from the individuals of the 7 populations with the mitochondrial genome sequence of *Eriocheir sinensis* as reference [88]. After assembly, the mitochondrial genome sequences were manually inspected and five represented sequences from each population were selected for phylogenetic tree construction using IQtree with 1,000 bootstrap [82].

### Species tree construction and divergence time estimation

Protein sequences of *Daphnia pulex*, *Daphnia magna*, *P. hawaiensis*, *Hyalella azteca*, *L. vannamei*, *Homarus americanus*, *Procambarus clarkii*, *Portunus_trituberculatus*, *Scylla paramamosain*, *E. sinensis*, *E. japanocia*, *E. hepuensis*, and *Drosophila melanogaster* were downloaded from the NCBI database except for *E. japanocia* and *E. hepuensis*. Species trees were constructed by IQtree using the 684 one-to-one single-copy orthologous genes identified by Orthofinder v2.3.11 software[89]. We further employed MCMCTree in PAML (version 4.8) [90] to estimate the divergence times of the species using the calibration time as a constraint, including *Drosophila melanogaster* – *Daphnia pulex* (474.8 - 530.0 million years ago), *Homarus americanus* – *Procambarus clarkii* (241.0 - 321.6 million years ago), and *Daphnia magna* – *Daphnia pulex* (130-158 million years ago) which were derived from the TimeTree database [91]. The MCMCTree was run for 500,000 interactions, and the first 50,000 samples were burned in. We ran the program three times for each data type to confirm that the results were similar between runs.

## Ethical statement

Our study, sampling procedures, and experimental protocols were approved by the Institutional Animal Care and Use Committee of Shanghai Ocean University (Shanghai, China) on the care and use of animals for scientific purposes.

## Data Availability

The raw resequencing data of the seven populations were deposited at the National Genomics Data Center, Beijing Institute of Genomics, Chinese Academy of Sciences / China National Center for Bioinformation (GSA ID: CRA013348). The RNA-seq data of *Eriocheir japonica* and *Eriocheir hepuensis* were available at https://ngdc.cncb.ac.cn/gsa/browse/CRA020935 (GSA ID: CRA020935).

## Authors’ contributions

**Jun Wang**: Conceptualization, Methodology, Data analysis, Visualization, Writing—original draft, Writing—review & editing. **Xin Hou**: Data curation, Visualization. Xiaowen Chen: Methodology, Data curation. **Roland Nathan Mandal**: Data analysis, Visualization. **Nusrat Hasan Kanika**: Visualization, Writing—original draft, Writing—review & editing. **Chunhong Yuan**: Sample collection, Writing—review & editing. **Yongju Luo**: Sample collection, Writing—review & editing. **Chenghui Wang**: Conceptualization, Methodology, Supervision, Writing—original draft, Writing—review & editing.

## Acknowledgments

This work was supported by the National Key Research and Development Project of China (Grant No. 2022YFD2400701).

