## Supplementary Figures 1-10 for "Genomic Insights into Hybridization and Speciation of Mitten Crabs in the *Eriocheir* Genus"

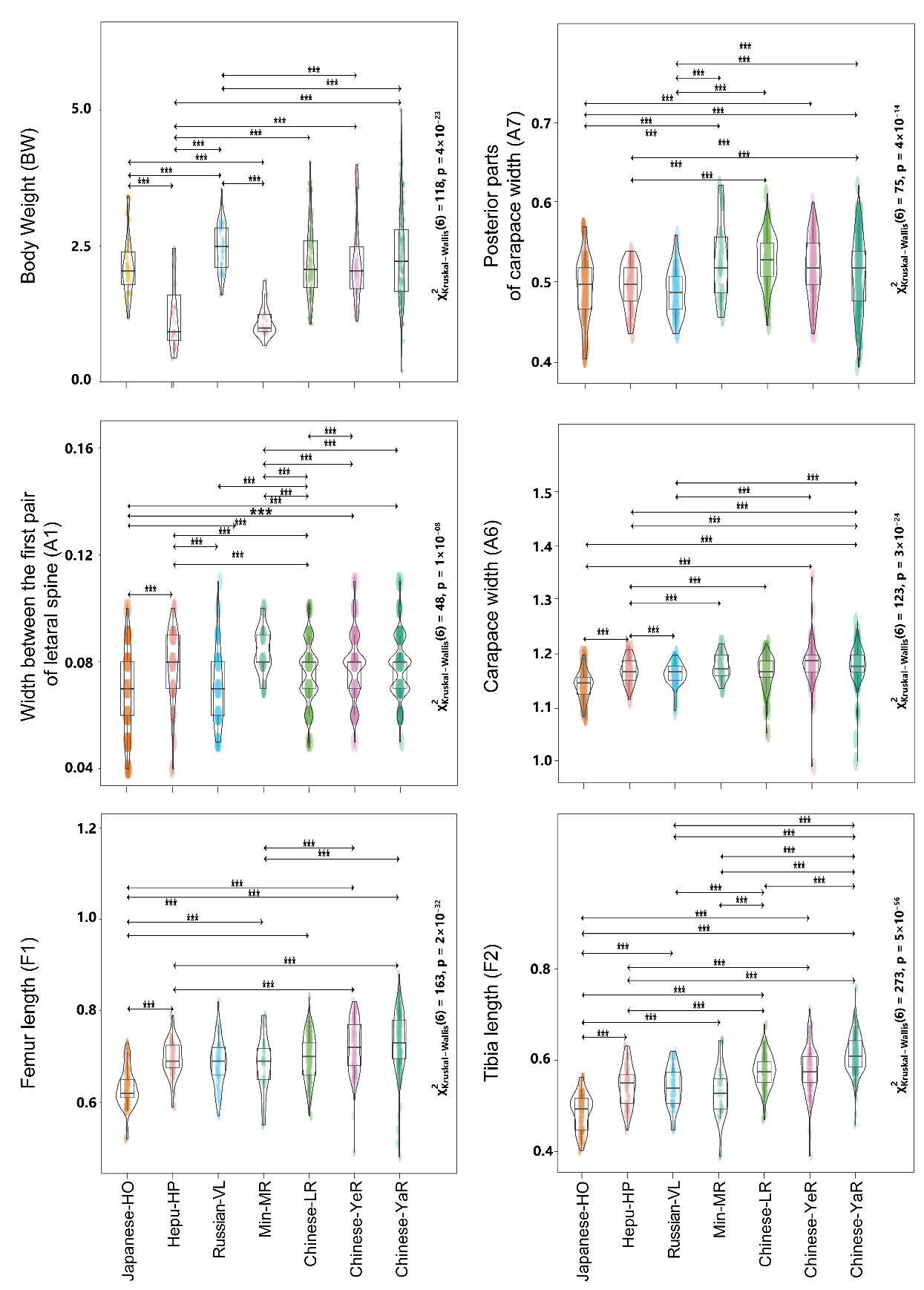

**Figure S1. Comparative Analysis of Morphological Traits in Seven Mitten Crab Populations Using ANOVA.**

Violin plots display the variation in phenotypic traits (A1, A6, A7, F1, F2) and body weight (BW) across mitten crab populations from Japanese-HO, Hepu-HP, Russian-VL, Min -MR, Chinese- YeR, YaR, LR. For consistency, all morphological traits (A1, A6, A7, F1, F2) were normalized by shell length (L), with the exception of body weight. Significant differences between groups are indicated by "***" (*p* < 0.001). Refer to Figure 1c for trait definitions.

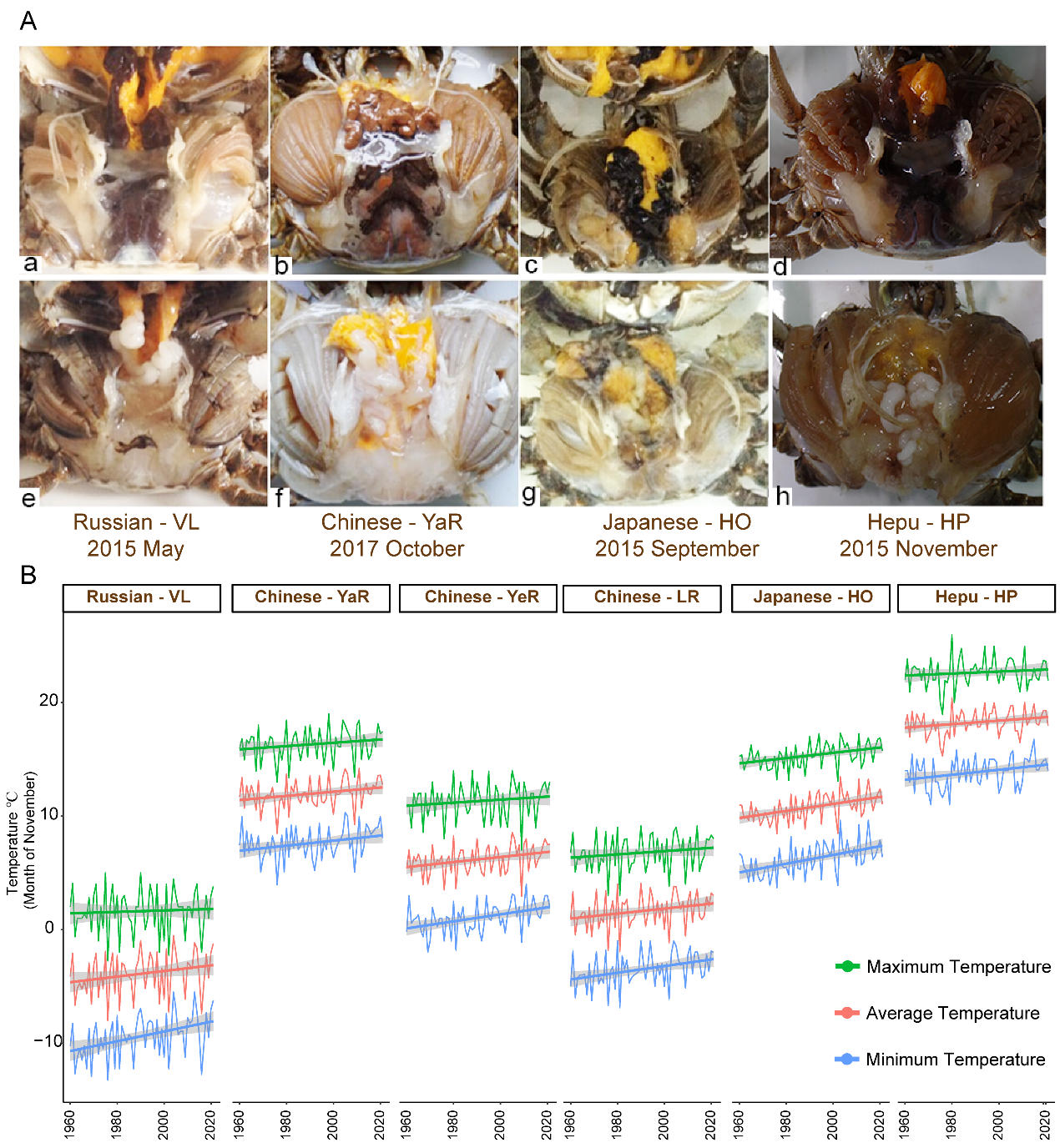

**Figure S2. Reproductive Maturity Seasons Across Mitten Crab Species from Varied Geographic Regions.** (A) Ovary and Testis Characteristics of Mitten Crab Species: (a, e) Russian mitten crab, (b, f) Chinese mitten crab, (c, g) Japanese mitten crab, and (d, h) Hepu mitten crab. (B) Graphs illustrate temperature changes in physiological indices across different geographic populations.

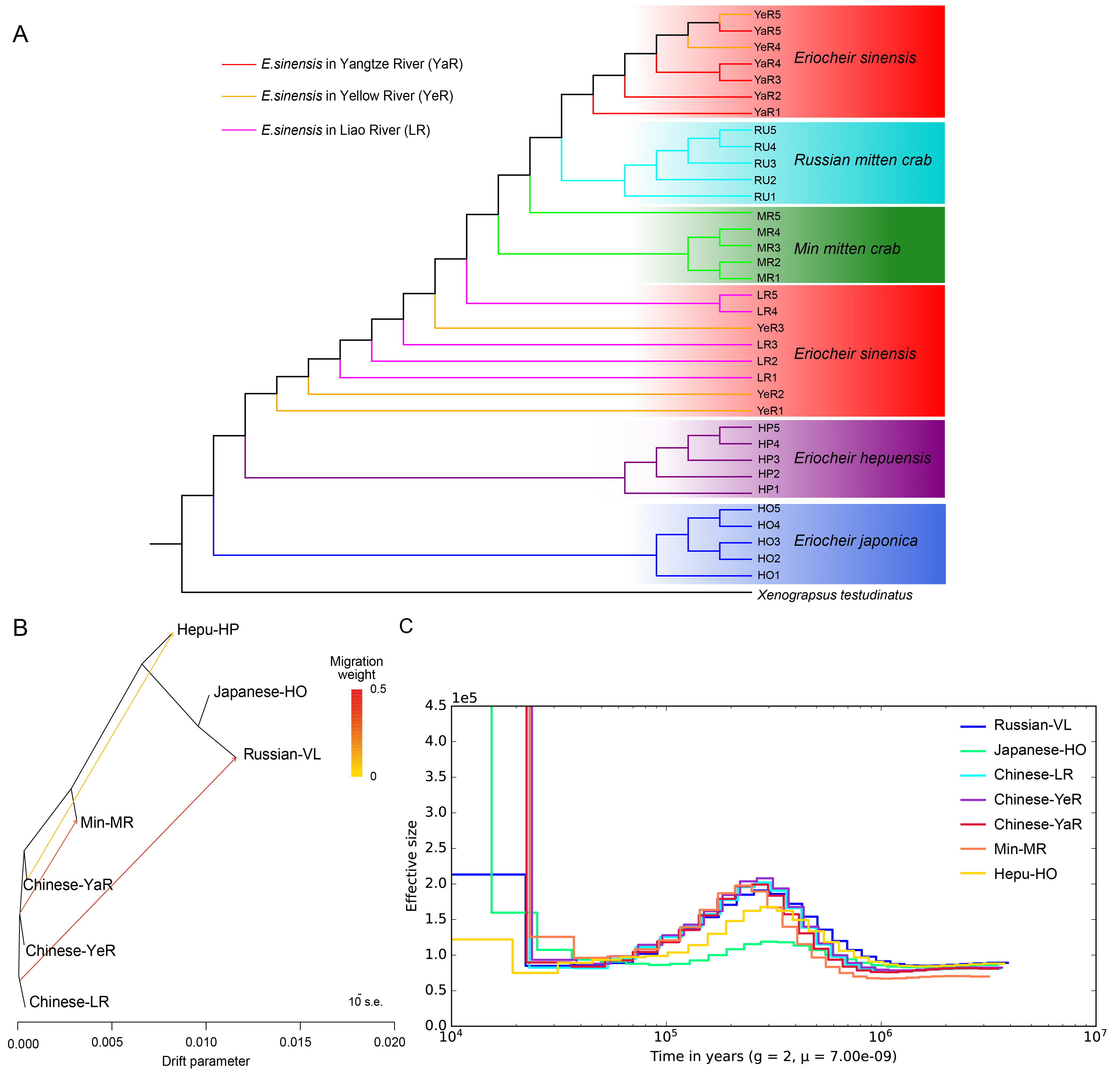

**Figure S3.** (A) Phylogenetic Tree of Mitochondrial Genomes in Mitten Crabs from the 7 Populations, with *Xenograpsus testudinatus* (NC_013480) Used as an Outgroup. (B) Treemix analysis displaying inferred migration edges among those populations. (C) Demographic history of mitten crabs from the 7 populations.

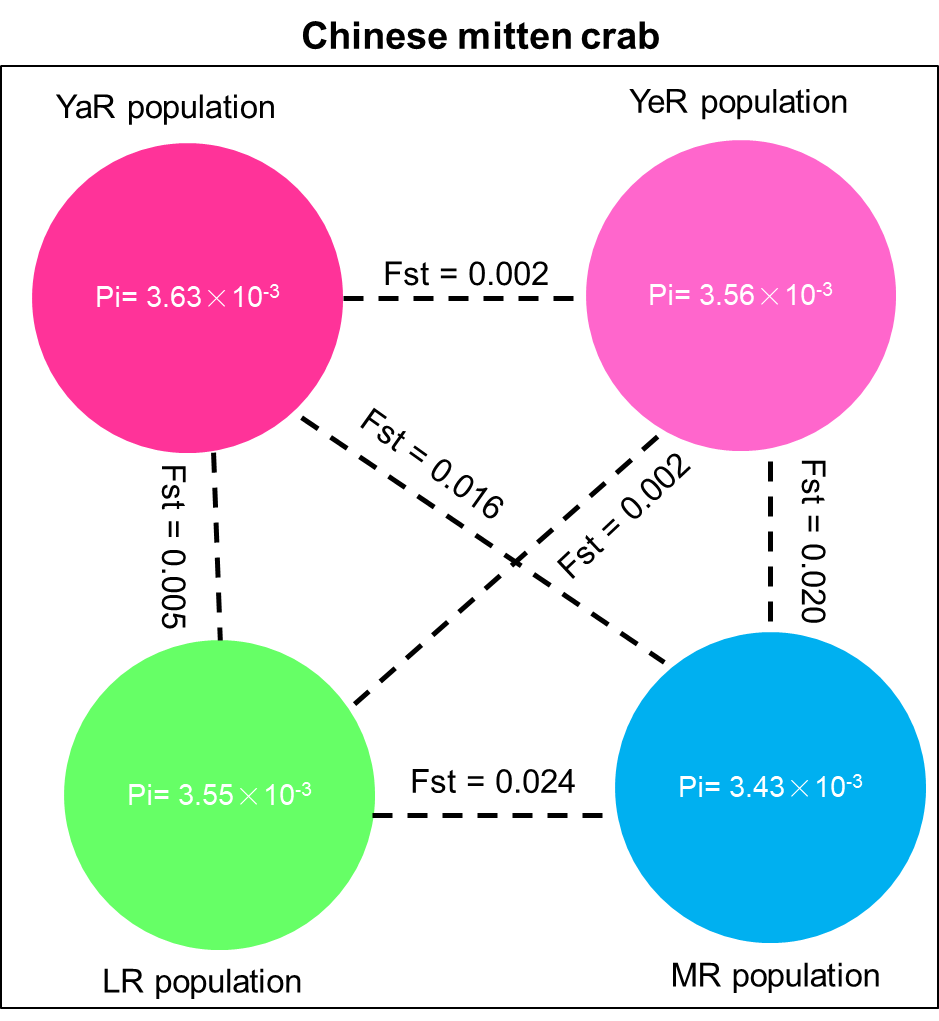

**Figure S4. Genetic Diversity and Genetic Differentiation Values Among Chinese Mitten Crab Populations.**

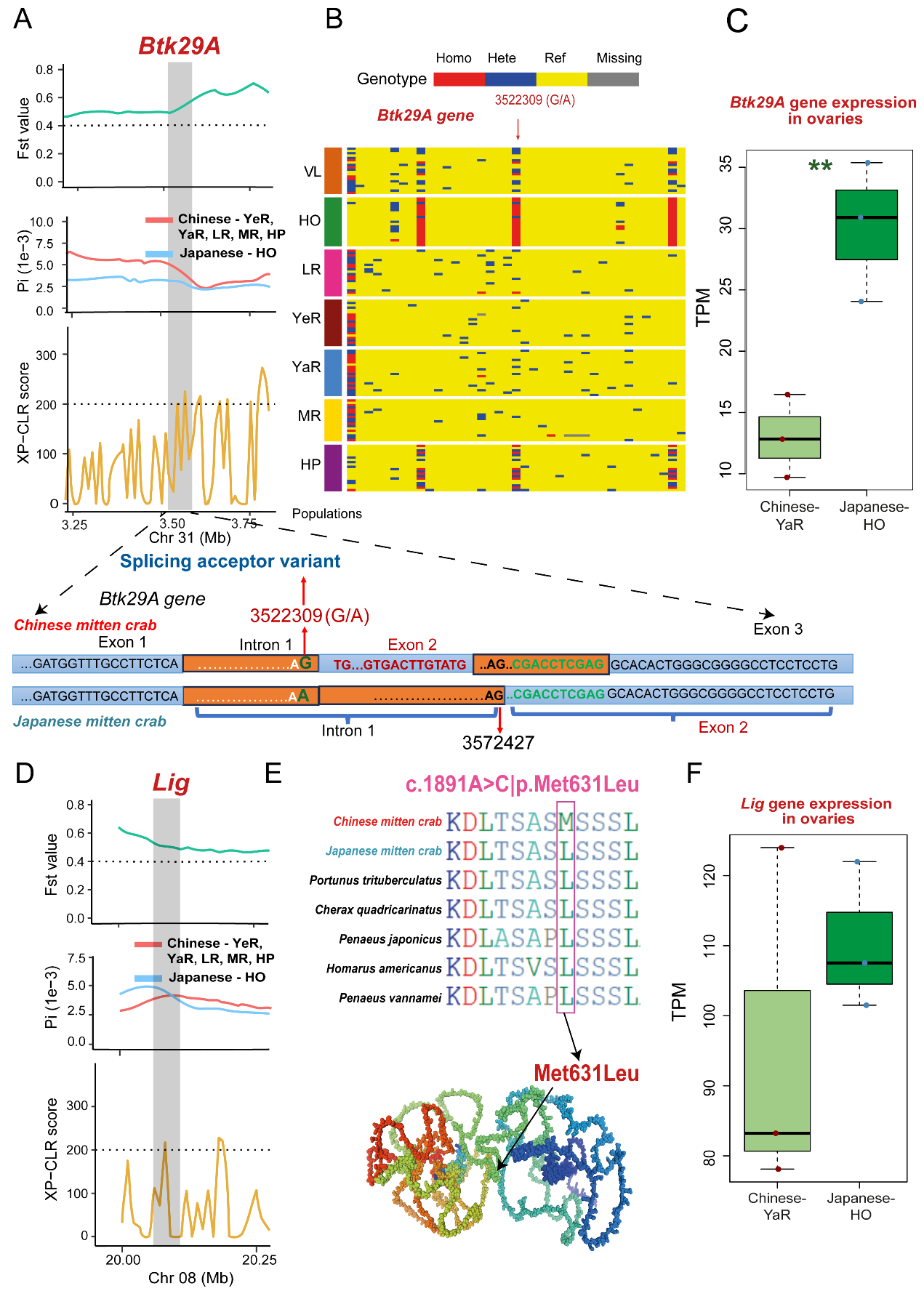

**Figure S5. Divergent selection between Chinese and Japanese mitten crab populations, based on *Btk29A* and *Lig* gene**.

(A) Detection of selection signals in the *Btk29A* gene, with a comparison of gene structure between Chinese and Japanese mitten crabs. (B) Haplotype diversity of the Btk29A gene across different mitten crab populations. (C) *Btk29A* gene expression in the ovary, demonstrating significant differential expression between Chinese-YaR and Japanese-HO mitten crabs (*p* <0.01). (D) Identification of selection signals in the *Lig* gene. (E) Sequence alignment of partial Lig proteins, highlighting mutations. (F) *Lig* gene expression in the ovary, showing distinct expression patterns between Chinese-YaR and Japanese-HO mitten crabs.

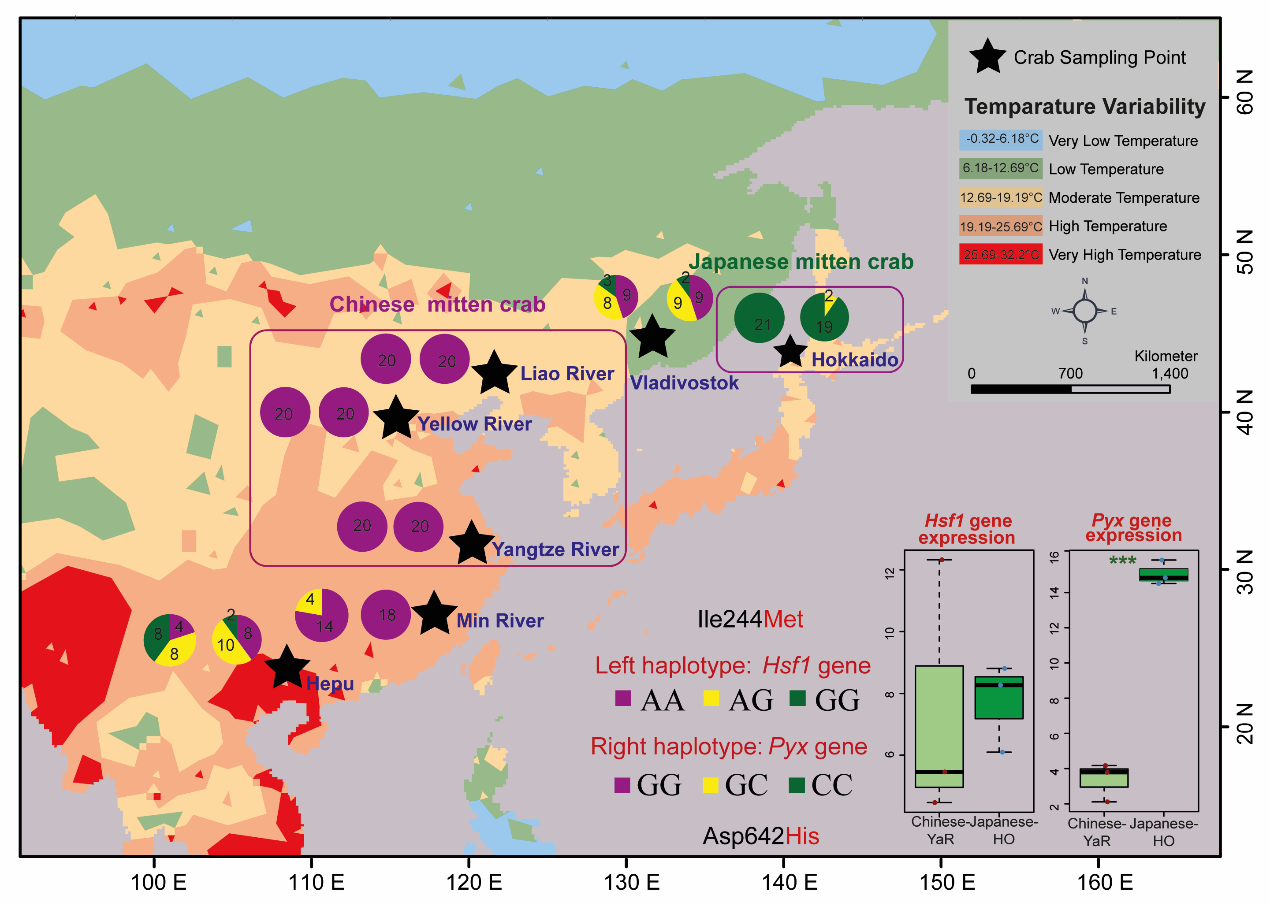

**Figure S6.** Observations of divergent selection in the *Hsf1* and *Pyx* genes between Chinese and Japanese mitten crabs. The two circle indicates the haplotype of *Hsf1* (left circle) and *Pyx* (right cricle) gene. Number in the circle indicates the number of individuals in this population with the specific genotype. The expression value (TPM) of *Hsf1* and *Pyx* gene in the ovary of Chinese and Japanese mitten crab were present in the right corner. The temperature data were downloaded from the WroldClim2 website (https://www.worldclim.org/data/worldclim21.html).

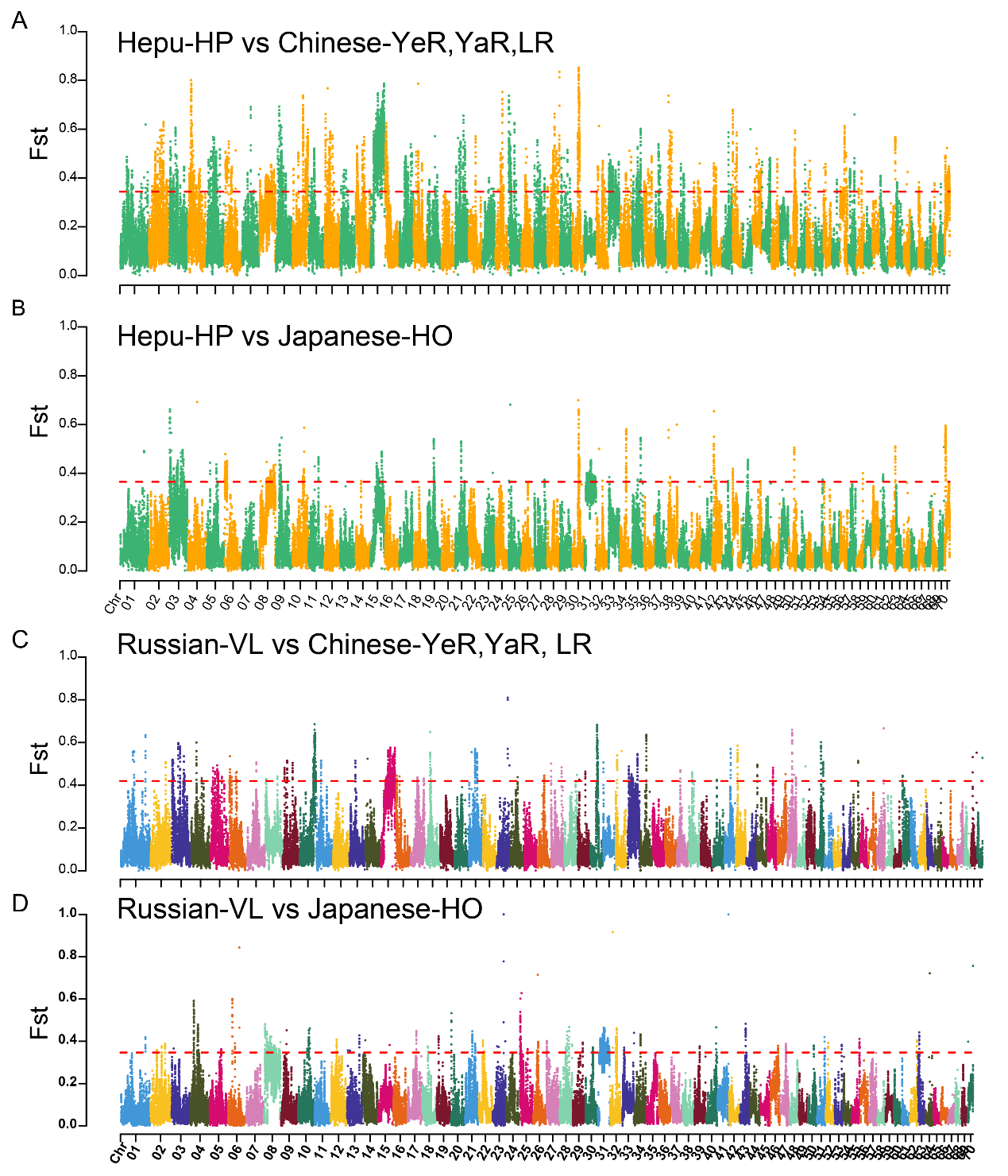

**Figure S7.** Genome-Wide Genetic Differentiation (*Fst* Value) Between (A) Hepu Mitten Crab and Chinese Mitten Crab and (B) Hepu Mitten Crab and Japanese Mitten Crabs. (C) Russian Mitten Crab and Chinese Mitten Crab and (D) Russian Mitten Crab and Japanese Mitten Crabs.

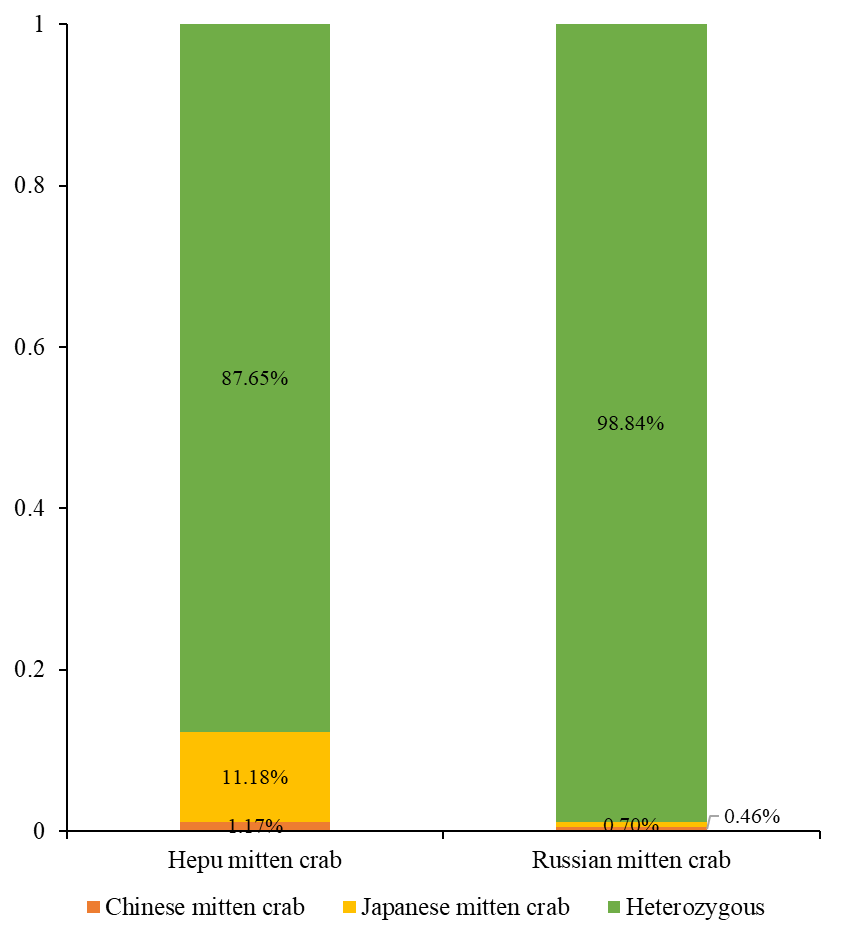

**Figure S8.** Frequency of Divergent Allele Genotypes from Chinese-YeR, YaR, LR, and Japanese-HO Mitten Crabs in Hepu-HP and Russian-VL Mitten Crabs.

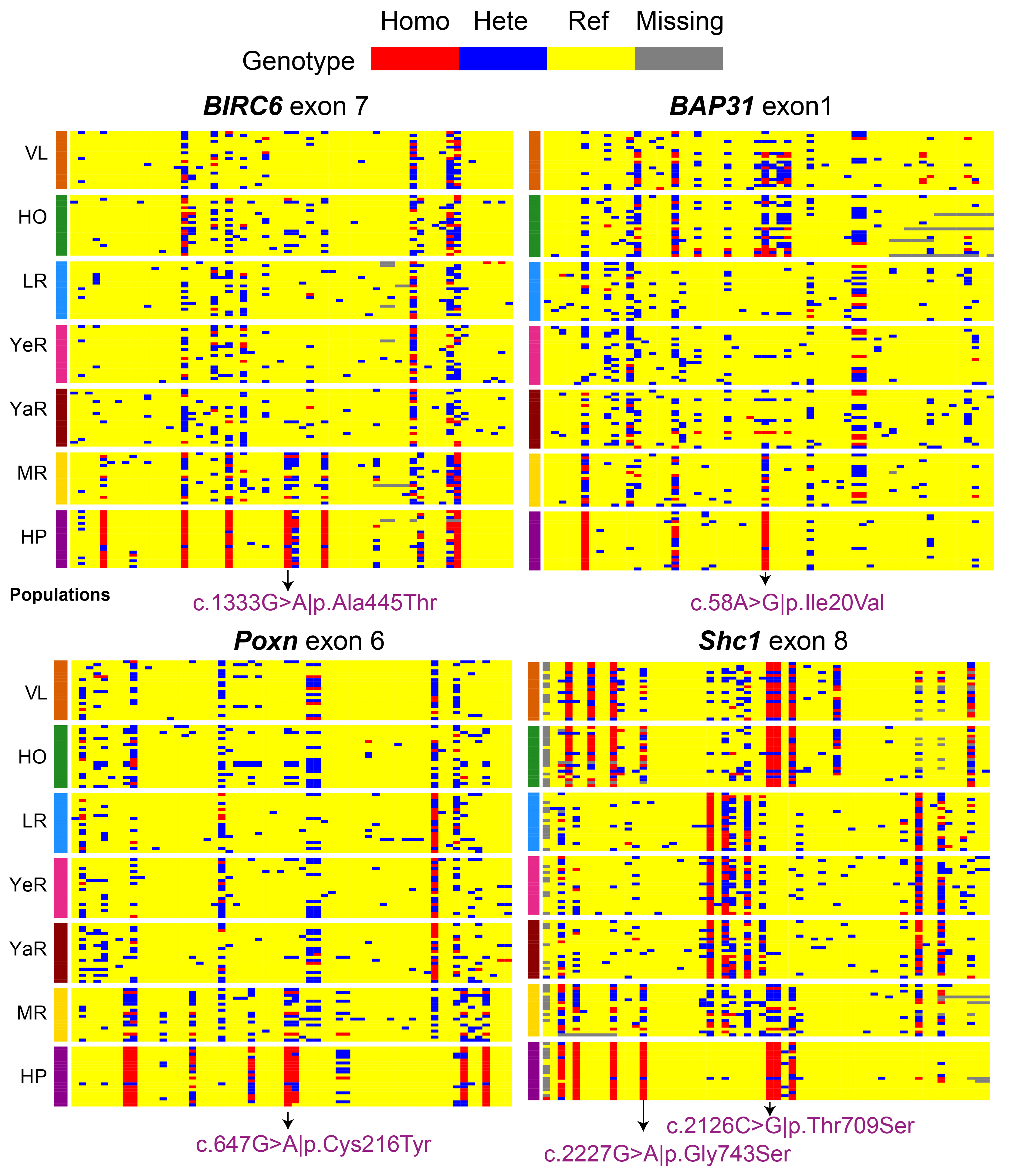

**Figure S9.** Haplotype Figures of *Birc6, Bap31, Poxn, and Shc1* genes among the selected populations of mitten crab.

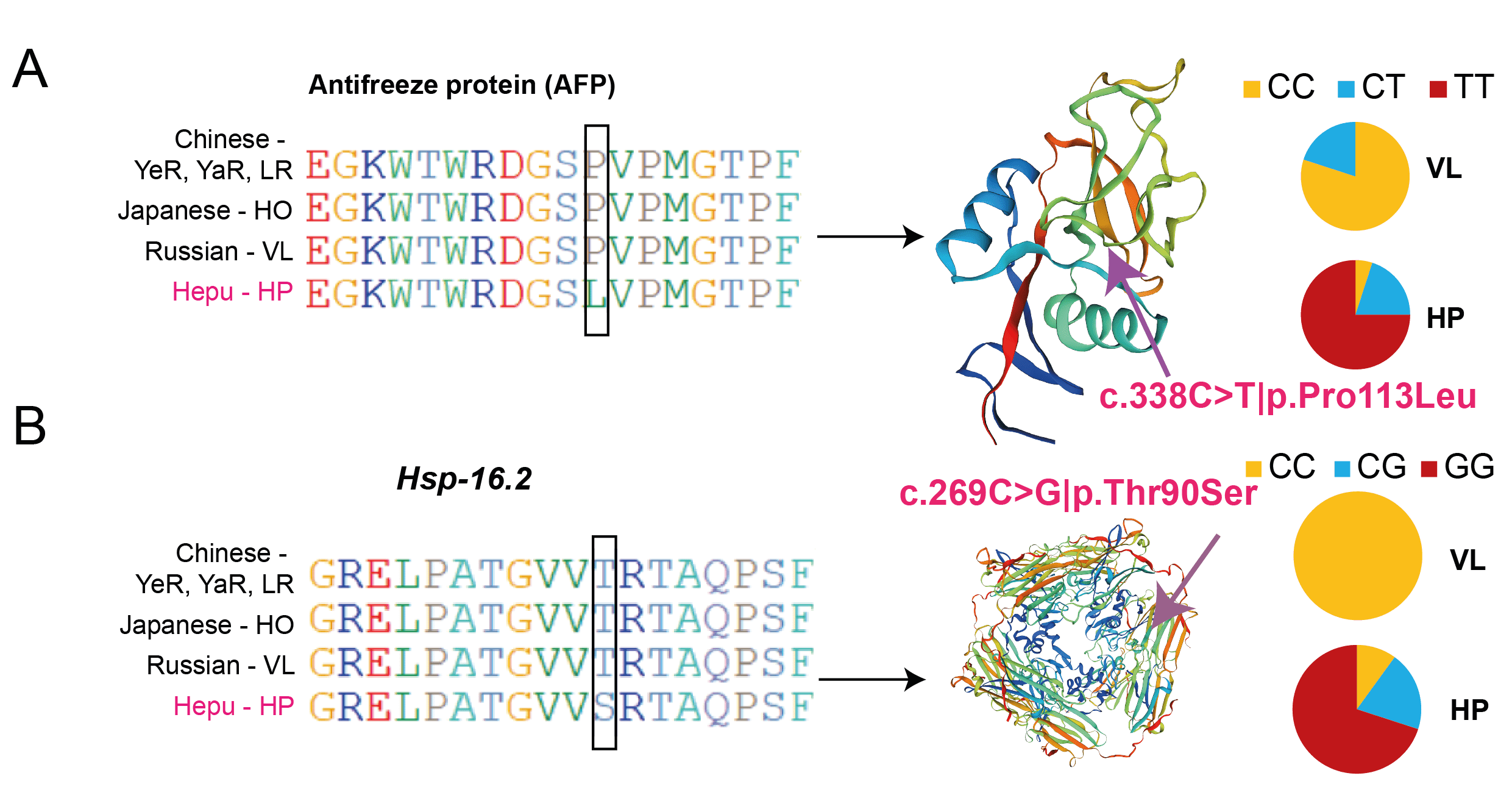

**Figure S10.** (A) Sequence alignment, protein structure, and haplotype frequency of the *AFP* gene. (B) Sequence alignment, protein structure, and haplotype frequency of the Hsp-16.2 gene.
