## Supplementary Tables 1-8 for "Genomic Insights into Hybridization and Speciation of Mitten Crabs in the *Eriocheir* Genus"

**Table S1. Raw sequencing data information for *Eriocheir japonica* genome assembly across multiple sequencing platforms.**

| Insert size | Sequencing  Platform | Mode | No. of Reads | No. of Bases | Sequencing Depth (X) |
| --- | --- | --- | --- | --- | --- |
| 180 | Illumina HiSeq 2500 | PE125 | 509,344,202 | 64,177,369,452 | 41.95 |
| 500 | Illumina HiSeq 2500 | PE125 | 445,488,362 | 56,131,533,612 | 36.69 |
| 800 | Illumina HiSeq 2500 | PE125 | 422,638,190 | 52,829,773,750 | 34.53 |
| 2k | Illumina HiSeq 2500 | PE125 | 503,451,454 | 63,434,883,204 | 41.46 |
| 5k | Illumina HiSeq 2500 | PE125 | 317,390,542 | 39,991,208,292 | 26.14 |
| 10k | Illumina HiSeq 2500 | PE125 | 126,462,706 | 15,934,300,956 | 10.41 |
| 10k | Illumina HiSeq 4000 | PE150 | 224,930,908 | 33,739,636,200 | 22.05 |
| Total | - | - | 2,549,706,364 | 326,238,705,466 | 213.23 |
|  | PacBio platform | - | 3,559,898 | 21,799,025,296 | 14.25 |
|  | 10X Genomics platform | PE150 | 784,212,882 | 117,631,932,300 | 76.88 |

Raw reads data generated for the *Eriocheir japonica* genome sequencing project using Illumina platform, PacBio platform and 10X genomics platforms.

**Table S2. Raw sequencing data information for *Eriocheir hepuensis* genome assembly.**

| Insert size | Sequencing Platform | Mode | No. of Reads | No. of Bases | Sequencing Depth (X) |
| --- | --- | --- | --- | --- | --- |
| 250 | Illumina HiSeq 4000 | PE150 | 823,719,318 | 123,557,897,700 | 76.27 |
| 400 | Illumina HiSeq 4000 | PE150 | 736,711,770 | 110,506,765,500 | 68.21 |
| 800 | Illumina HiSeq 4000 | PE150 | 458,664,708 | 57,333,088,500 | 35.39 |
| 2k | Illumina HiSeq 4000 | PE150 | 766,300,218 | 114,945,032,700 | 70.95 |
| 5k | Illumina HiSeq 4000 | PE150 | 466,914,348 | 70,037,152,200 | 43.23 |
| 10k | Illumina HiSeq 4000 | PE150 | 355,560,222 | 53,334,033,300 | 32.92 |
| Total |  |  | 3,607,870,58 | 529,713,969,900 | 326.98 |

**Table S3. Genome assembly metrics of three mitten crab species: *Eriocheir sinensis*, *Eriocheir japonica*, and *Eriocheir hepuensis*.**

|  | ***Eriocheir sinensis**** | ***Eriocheir japonica*** | ***Eriocheir hepuensis*** |
| --- | --- | --- | --- |
| **Genome assembly statistics** | | | |
| **Total assembled genome size (bp)** | 1,767,846,446 | 1,238,768,649 | 1,183,716,478 |
| **Total number of scaffolds** | 2,160 | 45,502 | 78,603 |
| **No. of scaffolds ≥1000 bp** | 2,158 | 23,785 | 40,046 |
| **No. of scaffolds ≥5000 bp** | 2,107 | 12,169 | 20,461 |
| **Longest length (bp)** | 45,379,147 | 3,836,756 | 1,105,787 |
| **Scaffold N50 (bp) / No. of N50** | 16,975,517 / 40 | 442,749/706 | 122,458/2,581 |
| **Contig N50 (bp) / No. of Contig N50** | 717,335 / 434 | 15,754/18,189 | 15,110/18,742 |
| **Genomic features** | | | |
| **GC (%)** | 41.21 | 42.73 | 43.29 |
| **Predicted protein-coding genes** | 20,286 | 18,418 | 19,253 |
| **Annotated genes** | 18,507 | 15,844 | 16,260 |

* The genome assembly data of *Eriocheir sinensis* were obtained from NCBI database, <https://www.ncbi.nlm.nih.gov/datasets/genome/GCF_024679095.1/>.

**Table S4.** **Comparison of BUSCO assessment results among *Eriocheir* genomes.**

| **Type** | ***E. sinensis****  **(Number/Percent)** | ***E. japonica***  **(Number/Percent)** | ***E. hepuensis***  **(Number/Percent)** |
| --- | --- | --- | --- |
| Complete BUSCOs (C) | 1009/94.65 | 956/89.68 | 934/87.62 |
| Complete and single-copy BUSCOs (S) | 942/88.37 | 911/85.46 | 913/85.65 |
| Complete and duplicated BUSCOs (D) | 67/6.29 | 45/4.22 | 21/1.97 |
| Fragmented BUSCOs (F) | 7/0.66 | 46/4.32 | 64/6.00 |
| Missing BUSCOs (M) | 50/4.69 | 64/6.00 | 68/6.38 |
| Total BUSCO groups searched | 1066/100 | 1066/100 | 1066/100 |

* The genome assembly data of *Eriocheir sinensis* were obtained from NCBI database, <https://www.ncbi.nlm.nih.gov/datasets/genome/GCF_024679095.1/>.

**Table S5. Information of sampling localities, and sample size of the 7 populations of *Eriocheir* species.**

| Location | Latitude | Longitude | Site Abbreviation | Sample Identity | Sample Size | Year of Collection |
| --- | --- | --- | --- | --- | --- | --- |
| Vladivostok, Russia | 43.2 | 131.9 | VL | Russian - VL | 32 | 2015 |
| Hokkaido, Japan | 43.1 | 141.6 | HO | Japanese - HO | 45 | 2015 |
| Liao River, China | 41.3 | 122.3 | LR | Chinese - LR | 52 | 2017 |
| Yellow River, China | 38.1 | 119.1 | YeR | Chinese - YeR | 54 | 2017 |
| Yangtze River, China | 31.6 | 121.6 | YaR | Chinese -YaR | 55 | 2017 |
| Min River, China | 26.0 | 119.3 | MR | Min - MR | 22 | 2015 |
| Hepu, China | 21.4 | 109.0 | HP | Hepu - HP | 29 | 2015 |

**Table S6. Sequencing and mapping information of the 139 mitten crab individuals from the 7 populations of *Eriocheir* species.**

| **Sample ID** | **Populations** | **Raw reads** | **Raw Data** | **Raw Depth** | **Clean reads** | **Clean data** | **Depth after filter** | **Mapping rate** |
| --- | --- | --- | --- | --- | --- | --- | --- | --- |
| YeR1 | YeR | 277,964,696 | 41,694,704,400 | 25 | 269,381,570 | 40,407,235,500 | 24 | 0.99 |
| YeR2 | YeR | 261,145,332 | 39,171,799,800 | 23 | 252,866,914 | 37,930,037,100 | 22 | 0.99 |
| YeR3 | YeR | 262,955,878 | 39,443,381,700 | 23 | 254,689,204 | 38,203,380,600 | 22 | 0.99 |
| YeR4 | YeR | 301,618,624 | 45,242,793,600 | 27 | 292,669,422 | 43,900,413,300 | 26 | 0.99 |
| YeR5 | YeR | 250,209,592 | 37,531,438,800 | 22 | 225,308,262 | 33,796,239,300 | 20 | 0.98 |
| YeR6 | YeR | 249,510,936 | 37,426,640,400 | 22 | 222,783,192 | 33,417,478,800 | 20 | 0.98 |
| YeR7 | YeR | 242,802,646 | 36,420,396,900 | 21 | 234,921,442 | 35,238,216,300 | 21 | 0.99 |
| YeR8 | YeR | 250,846,576 | 37,626,986,400 | 22 | 231,022,780 | 34,653,417,000 | 20 | 0.99 |
| YeR9 | YeR | 252,456,360 | 37,868,454,000 | 22 | 224,185,674 | 33,627,851,100 | 20 | 0.99 |
| YeR10 | YeR | 252,465,796 | 37,869,869,400 | 22 | 244,234,774 | 36,635,216,100 | 22 | 0.98 |
| YeR11 | YeR | 257,752,936 | 38,662,940,400 | 23 | 249,892,852 | 37,483,927,800 | 22 | 0.99 |
| YeR12 | YeR | 244,588,786 | 36,688,317,900 | 22 | 237,207,072 | 35,581,060,800 | 21 | 0.96 |
| YeR13 | YeR | 262,555,878 | 39,383,381,700 | 23 | 254,397,988 | 38,159,698,200 | 22 | 0.97 |
| YeR14 | YeR | 250,086,152 | 37,512,922,800 | 22 | 221,353,204 | 33,202,980,600 | 20 | 0.97 |
| YeR15 | YeR | 240,842,158 | 36,126,323,700 | 21 | 233,213,300 | 34,981,995,000 | 21 | 0.98 |
| YeR16 | YeR | 253,015,064 | 37,952,259,600 | 22 | 226,907,030 | 34,036,054,500 | 20 | 0.90 |
| YeR17 | YeR | 240,173,074 | 36,025,961,100 | 21 | 232,192,692 | 34,828,903,800 | 20 | 0.99 |
| YeR18 | YeR | 254,158,734 | 38,123,810,100 | 22 | 246,423,412 | 36,963,511,800 | 22 | 0.99 |
| YeR19 | YeR | 261,581,424 | 39,237,213,600 | 23 | 252,904,620 | 37,935,693,000 | 22 | 0.99 |
| YeR20 | YeR | 252,128,386 | 37,819,257,900 | 22 | 244,729,130 | 36,709,369,500 | 22 | 0.99 |
| YaR1 | YaR | 270,255,532 | 40,538,329,800 | 24 | 261,935,878 | 39,290,381,700 | 23 | 0.99 |
| YaR2 | YaR | 307,490,860 | 46,123,629,000 | 27 | 297,576,038 | 44,636,405,700 | 26 | 0.97 |
| YaR3 | YaR | 263,761,524 | 39,564,228,600 | 23 | 255,782,928 | 38,367,439,200 | 23 | 0.98 |
| YaR4 | YaR | 261,584,060 | 39,237,609,000 | 23 | 253,648,076 | 38,047,211,400 | 22 | 0.99 |
| YaR5 | YaR | 259,457,574 | 38,918,636,100 | 23 | 251,064,690 | 37,659,703,500 | 22 | 0.99 |
| YaR6 | YaR | 244,102,860 | 36,615,429,000 | 22 | 236,662,958 | 35,499,443,700 | 21 | 0.98 |
| YaR7 | YaR | 247,220,518 | 37,083,077,700 | 22 | 238,796,120 | 35,819,418,000 | 21 | 0.98 |
| YaR8 | YaR | 239,894,214 | 35,984,132,100 | 21 | 232,440,936 | 34,866,140,400 | 21 | 0.97 |
| YaR9 | YaR | 252,106,006 | 37,815,900,900 | 22 | 244,630,980 | 36,694,647,000 | 22 | 0.99 |
| YaR10 | YaR | 254,999,724 | 38,249,958,600 | 22 | 231,069,666 | 34,660,449,900 | 20 | 0.98 |
| YaR11 | YaR | 249,461,166 | 37,419,174,900 | 22 | 241,404,986 | 36,210,747,900 | 21 | 0.99 |
| YaR12 | YaR | 253,567,652 | 38,035,147,800 | 22 | 245,554,942 | 36,833,241,300 | 22 | 0.98 |
| YaR13 | YaR | 270,633,608 | 40,595,041,200 | 24 | 262,815,086 | 39,422,262,900 | 23 | 0.99 |
| YaR14 | YaR | 303,882,160 | 45,582,324,000 | 27 | 294,046,740 | 44,107,011,000 | 26 | 0.99 |
| YaR15 | YaR | 297,240,896 | 44,586,134,400 | 26 | 288,448,846 | 43,267,326,900 | 25 | 0.98 |
| YaR16 | YaR | 272,490,496 | 40,873,574,400 | 24 | 264,241,812 | 39,636,271,800 | 23 | 0.99 |
| YaR17 | YaR | 241,169,576 | 36,175,436,400 | 21 | 233,705,024 | 35,055,753,600 | 21 | 0.99 |
| YaR18 | YaR | 307,578,900 | 46,136,835,000 | 27 | 297,906,332 | 44,685,949,800 | 26 | 0.99 |
| YaR19 | YaR | 252,879,692 | 37,931,953,800 | 22 | 219,598,032 | 32,939,704,800 | 19 | 0.98 |
| YaR20 | YaR | 255,472,268 | 38,320,840,200 | 23 | 247,951,680 | 37,192,752,000 | 22 | 0.99 |
| LR1 | LR | 277,387,740 | 41,608,161,000 | 24 | 268,623,922 | 40,293,588,300 | 24 | 0.98 |
| LR2 | LR | 244,354,130 | 36,653,119,500 | 22 | 236,510,272 | 35,476,540,800 | 21 | 0.97 |
| LR3 | LR | 267,048,732 | 40,057,309,800 | 24 | 258,605,596 | 38,790,839,400 | 23 | 0.94 |
| LR4 | LR | 264,376,432 | 39,656,464,800 | 23 | 255,729,958 | 38,359,493,700 | 23 | 0.99 |
| LR5 | LR | 255,845,530 | 38,376,829,500 | 23 | 247,672,560 | 37,150,884,000 | 22 | 0.99 |
| LR6 | LR | 251,469,764 | 37,720,464,600 | 22 | 223,550,068 | 33,532,510,200 | 20 | 0.98 |
| LR7 | LR | 288,335,486 | 43,250,322,900 | 25 | 279,109,218 | 41,866,382,700 | 25 | 0.99 |
| LR8 | LR | 262,263,796 | 39,339,569,400 | 23 | 254,071,882 | 38,110,782,300 | 22 | 0.98 |
| LR9 | LR | 256,666,272 | 38,499,940,800 | 23 | 248,619,408 | 37,292,911,200 | 22 | 0.99 |
| LR10 | LR | 280,170,930 | 42,025,639,500 | 25 | 271,953,920 | 40,793,088,000 | 24 | 0.98 |
| LR11 | LR | 275,127,936 | 41,269,190,400 | 24 | 266,423,910 | 39,963,586,500 | 24 | 0.98 |
| LR12 | LR | 267,866,446 | 40,179,966,900 | 24 | 259,455,548 | 38,918,332,200 | 23 | 0.98 |
| LR13 | LR | 262,899,932 | 39,434,989,800 | 23 | 254,387,652 | 38,158,147,800 | 22 | 0.99 |
| LR14 | LR | 276,885,246 | 41,532,786,900 | 24 | 268,378,148 | 40,256,722,200 | 24 | 0.99 |
| LR15 | LR | 248,258,694 | 37,238,804,100 | 22 | 240,274,580 | 36,041,187,000 | 21 | 0.99 |
| LR16 | LR | 267,323,106 | 40,098,465,900 | 24 | 258,354,810 | 38,753,221,500 | 23 | 0.99 |
| LR17 | LR | 272,872,468 | 40,930,870,200 | 24 | 263,752,182 | 39,562,827,300 | 23 | 0.98 |
| LR18 | LR | 253,270,920 | 37,990,638,000 | 22 | 245,179,362 | 36,776,904,300 | 22 | 0.98 |
| LR19 | LR | 247,655,062 | 37,148,259,300 | 22 | 206,582,736 | 30,987,410,400 | 18 | 0.98 |
| LR20 | LR | 261,069,530 | 39,160,429,500 | 23 | 252,748,058 | 37,912,208,700 | 22 | 0.98 |
| HP1 | HP | 236,369,296 | 35,455,394,400 | 21 | 224,026,744 | 33,604,011,600 | 20 | 0.99 |
| HP2 | HP | 286,707,246 | 43,006,086,900 | 25 | 263,196,384 | 39,479,457,600 | 23 | 0.99 |
| HP3 | HP | 263,403,988 | 39,510,598,200 | 23 | 243,259,012 | 36,488,851,800 | 21 | 0.99 |
| HP4 | HP | 287,213,280 | 43,081,992,000 | 25 | 263,427,980 | 39,514,197,000 | 23 | 0.99 |
| HP5 | HP | 262,735,988 | 39,410,398,200 | 23 | 238,842,130 | 35,826,319,500 | 21 | 0.99 |
| HP6 | HP | 221,539,760 | 33,230,964,000 | 20 | 192,263,198 | 28,839,479,700 | 17 | 0.99 |
| HP7 | HP | 263,069,742 | 39,460,461,300 | 23 | 237,003,546 | 35,550,531,900 | 21 | 0.99 |
| HP8 | HP | 226,709,424 | 34,006,413,600 | 20 | 202,516,424 | 30,377,463,600 | 18 | 0.99 |
| HP9 | HP | 243,467,922 | 36,520,188,300 | 21 | 243,781,782 | 36,567,267,300 | 22 | 0.99 |
| HP10 | HP | 263,842,892 | 39,576,433,800 | 23 | 243,781,782 | 36,567,267,300 | 22 | 0.99 |
| HP11 | HP | 220,422,092 | 33,063,313,800 | 19 | 204,348,134 | 30,652,220,100 | 18 | 0.99 |
| HP12 | HP | 253,001,748 | 37,950,262,200 | 22 | 232,412,900 | 34,861,935,000 | 21 | 0.99 |
| HP13 | HP | 229,662,002 | 34,449,300,300 | 20 | 203,625,664 | 30,543,849,600 | 18 | 0.99 |
| HP14 | HP | 266,956,656 | 40,043,498,400 | 24 | 244,074,056 | 36,611,108,400 | 22 | 0.99 |
| HP15 | HP | 242,129,312 | 36,319,396,800 | 21 | 222,734,724 | 33,410,208,600 | 20 | 0.99 |
| HP16 | HP | 281,265,716 | 42,189,857,400 | 25 | 256,932,314 | 38,539,847,100 | 23 | 0.98 |
| HP17 | HP | 234,668,120 | 35,200,218,000 | 21 | 216,994,812 | 32,549,221,800 | 19 | 0.99 |
| HP18 | HP | 264,651,960 | 39,697,794,000 | 23 | 246,269,570 | 36,940,435,500 | 22 | 0.99 |
| HP19 | HP | 239,999,116 | 35,999,867,400 | 21 | 205,571,840 | 30,835,776,000 | 18 | 0.99 |
| HP20 | HP | 280,753,920 | 42,113,088,000 | 25 | 250,366,382 | 37,554,957,300 | 22 | 0.99 |
| HO1 | HO | 279,883,034 | 41,982,455,100 | 25 | 266,904,492 | 40,035,673,800 | 24 | 0.99 |
| HO2 | HO | 269,629,978 | 40,444,496,700 | 24 | 257,989,304 | 38,698,395,600 | 23 | 0.98 |
| HO3 | HO | 295,509,190 | 44,326,378,500 | 26 | 279,964,808 | 41,994,721,200 | 25 | 0.99 |
| HO4 | HO | 257,930,036 | 38,689,505,400 | 23 | 243,395,438 | 36,509,315,700 | 21 | 0.99 |
| HO5 | HO | 276,742,260 | 41,511,339,000 | 24 | 263,307,950 | 39,496,192,500 | 23 | 0.98 |
| HO6 | HO | 280,443,500 | 42,066,525,000 | 25 | 269,237,070 | 40,385,560,500 | 24 | 0.97 |
| HO7 | HO | 287,093,334 | 43,064,000,100 | 25 | 275,296,400 | 41,294,460,000 | 24 | 0.99 |
| HO8 | HO | 276,313,120 | 41,446,968,000 | 24 | 263,761,064 | 39,564,159,600 | 23 | 0.99 |
| HO9 | HO | 295,387,934 | 44,308,190,100 | 26 | 281,329,644 | 42,199,446,600 | 25 | 0.99 |
| HO10 | HO | 254,764,768 | 38,214,715,200 | 22 | 244,038,062 | 36,605,709,300 | 22 | 0.95 |
| HO11 | HO | 275,035,902 | 41,255,385,300 | 24 | 261,161,226 | 39,174,183,900 | 23 | 0.99 |
| HO12 | HO | 257,512,336 | 38,626,850,400 | 23 | 244,251,182 | 36,637,677,300 | 22 | 0.95 |
| HO13 | HO | 227,415,444 | 34,112,316,600 | 20 | 219,457,506 | 32,918,625,900 | 19 | 0.92 |
| HO14 | HO | 281,805,180 | 42,270,777,000 | 25 | 271,881,010 | 40,782,151,500 | 24 | 0.88 |
| HO15 | HO | 287,294,072 | 43,094,110,800 | 25 | 275,354,006 | 41,303,100,900 | 24 | 0.99 |
| HO16 | HO | 259,292,974 | 38,893,946,100 | 23 | 245,297,392 | 36,794,608,800 | 22 | 0.98 |
| HO17 | HO | 259,691,430 | 38,953,714,500 | 23 | 242,659,984 | 36,398,997,600 | 21 | 0.99 |
| HO18 | HO | 283,799,552 | 42,569,932,800 | 25 | 269,058,846 | 40,358,826,900 | 24 | 0.98 |
| HO19 | HO | 236,131,906 | 35,419,785,900 | 21 | 225,719,386 | 33,857,907,900 | 20 | 0.99 |
| HO20 | HO | 247,030,372 | 37,054,555,800 | 22 | 237,431,082 | 35,614,662,300 | 21 | 0.99 |
| HO21 | HO | 292,022,892 | 43,803,433,800 | 26 | 279,259,976 | 41,888,996,400 | 25 | 0.99 |
| MR1 | MR | 242,891,938 | 36,433,790,700 | 21 | 229,904,024 | 34,485,603,600 | 20 | 0.60 |
| MR2 | MR | 262,628,076 | 39,394,211,400 | 23 | 245,055,816 | 36,758,372,400 | 22 | 0.99 |
| MR3 | MR | 258,971,532 | 38,845,729,800 | 23 | 246,632,194 | 36,994,829,100 | 22 | 0.47 |
| MR4 | MR | 279,213,404 | 41,882,010,600 | 25 | 255,357,264 | 38,303,589,600 | 23 | 0.99 |
| MR5 | MR | 281,258,190 | 42,188,728,500 | 25 | 258,407,686 | 38,761,152,900 | 23 | 0.94 |
| MR6 | MR | 239,216,588 | 35,882,488,200 | 21 | 219,073,624 | 32,861,043,600 | 19 | 0.99 |
| MR7 | MR | 231,759,860 | 34,763,979,000 | 20 | 217,790,896 | 32,668,634,400 | 19 | 0.73 |
| MR8 | MR | 269,148,216 | 40,372,232,400 | 24 | 251,386,270 | 37,707,940,500 | 22 | 0.99 |
| MR9 | MR | 283,655,710 | 42,548,356,500 | 25 | 261,191,688 | 39,178,753,200 | 23 | 0.84 |
| MR10 | MR | 226,293,612 | 33,944,041,800 | 20 | 205,371,238 | 30,805,685,700 | 18 | 0.99 |
| MR11 | MR | 259,000,438 | 38,850,065,700 | 23 | 239,844,378 | 35,976,656,700 | 21 | 0.99 |
| MR12 | MR | 268,531,400 | 40,279,710,000 | 24 | 243,058,744 | 36,458,811,600 | 21 | 0.74 |
| MR13 | MR | 240,073,462 | 36,011,019,300 | 21 | 218,725,040 | 32,808,756,000 | 19 | 0.99 |
| MR14 | MR | 258,304,576 | 38,745,686,400 | 23 | 236,221,916 | 35,433,287,400 | 21 | 0.99 |
| MR15 | MR | 274,595,442 | 41,189,316,300 | 24 | 254,713,064 | 38,206,959,600 | 22 | 0.99 |
| MR16 | MR | 308,364,632 | 46,254,694,800 | 27 | 273,702,288 | 41,055,343,200 | 24 | 0.98 |
| MR17 | MR | 292,825,140 | 43,923,771,000 | 26 | 270,985,858 | 40,647,878,700 | 24 | 0.99 |
| MR18 | MR | 273,687,182 | 41,053,077,300 | 24 | 254,658,586 | 38,198,787,900 | 22 | 0.99 |
| VL1 | VL | 287,395,730 | 43,109,359,500 | 25 | 269,493,790 | 40,424,068,500 | 24 | 0.99 |
| VL2 | VL | 268,837,564 | 40,325,634,600 | 24 | 252,241,510 | 37,836,226,500 | 22 | 0.99 |
| VL3 | VL | 289,834,954 | 43,475,243,100 | 26 | 269,646,504 | 40,446,975,600 | 24 | 0.99 |
| VL4 | VL | 279,165,022 | 41,874,753,300 | 25 | 260,187,488 | 39,028,123,200 | 23 | 0.99 |
| VL5 | VL | 258,581,710 | 38,787,256,500 | 23 | 241,118,108 | 36,167,716,200 | 21 | 0.99 |
| VL6 | VL | 236,427,692 | 35,464,153,800 | 21 | 223,216,608 | 33,482,491,200 | 20 | 0.99 |
| VL7 | VL | 284,524,882 | 42,678,732,300 | 25 | 267,821,290 | 40,173,193,500 | 24 | 0.99 |
| VL8 | VL | 267,765,998 | 40,164,899,700 | 24 | 251,041,464 | 37,656,219,600 | 22 | 0.99 |
| VL9 | VL | 247,307,918 | 37,096,187,700 | 22 | 233,139,286 | 34,970,892,900 | 21 | 0.99 |
| VL10 | VL | 272,203,294 | 40,830,494,100 | 24 | 255,289,868 | 38,293,480,200 | 23 | 0.99 |
| VL11 | VL | 276,218,916 | 41,432,837,400 | 24 | 259,226,324 | 38,883,948,600 | 23 | 0.99 |
| VL12 | VL | 289,293,844 | 43,394,076,600 | 26 | 271,990,510 | 40,798,576,500 | 24 | 0.98 |
| VL13 | VL | 271,756,438 | 40,763,465,700 | 24 | 255,819,542 | 38,372,931,300 | 23 | 0.99 |
| VL14 | VL | 282,973,278 | 42,445,991,700 | 25 | 263,784,592 | 39,567,688,800 | 23 | 0.99 |
| VL15 | VL | 282,882,610 | 42,432,391,500 | 25 | 264,515,430 | 39,677,314,500 | 23 | 0.99 |
| VL16 | VL | 270,120,074 | 40,518,011,100 | 24 | 253,969,698 | 38,095,454,700 | 22 | 0.99 |
| VL17 | VL | 291,074,590 | 43,661,188,500 | 26 | 271,708,694 | 40,756,304,100 | 24 | 0.99 |
| VL18 | VL | 280,595,308 | 42,089,296,200 | 25 | 263,519,942 | 39,527,991,300 | 23 | 0.98 |
| VL19 | VL | 281,054,726 | 42,158,208,900 | 25 | 263,801,880 | 39,570,282,000 | 23 | 0.99 |
| VL20 | VL | 242,097,256 | 36,314,588,400 | 21 | 219,797,202 | 32,969,580,300 | 19 | 0.99 |
| Average |  | 263,777,605 | 39,566,640,714 | 23 | 248,605,364 | 37,290,804,587 | 22 | 0.97 |

**Table S7. SNPs annotation information by Snpeff software.**

| **Type** | **Count** | **Percentage** |
| --- | --- | --- |
| **Downstream** | 6,708,391 | 5.764% |
| **Exon** | 1,317,392 | 1.132% |
| **missense_variant** | 434,715 | 0.373% |
| **synonymous_variant** | 887,364 | 0.762% |
| **stop_gained** | 8,571 | 0.007% |
| **stop_lost** | 1,076 | 0.001% |
| **start_lost** | 1,040 | 0.001% |
| **Intergenic** | 80,318,875 | 69.014% |
| **Intron** | 21,033,433 | 18.073% |
| **Splice_Site_Acceptor** | 1,195 | 0.001% |
| **Splice_Site_Donor** | 1,935 | 0.002% |
| **Splice_Site_Region** | 109,853 | 0.094% |
| **Upstream** | 6,889,109 | 5.919% |
| **UTR_5_Prime** | 41 | 0% |

**Table S8. Pairwise Dxy values identified among different *Eriocheir* populations**

| Comparison Groups | Dxy value (10^-2^) |
| --- | --- |
| Chinese-YaR, YeR, LR vs Japanese-HO | 8.02 |
| Chinese-YaR, YeR, LR vs Hepu-HP | 7.55 |
| Chinese-YaR, YeR, LR vs Russian-VL | 7.37 |
| Japanese-HO vs Hepu-HP | 6.81 |
| Japanese-HO vs Russian-VL | 6.93 |
| Hepu-HP vs Russian-VL | 7.17 |
